# A Comparative Study of Topology-based Pathway Enrichment Analysis Methods

**DOI:** 10.1101/814145

**Authors:** Jing Ma, Ali Shojaie, George Michailidis

## Abstract

**Background:** Pathway enrichment analysis is extensively used in the analysis of Omics data for gaining biological insights into the functional roles of pre-defined subsets of genes, proteins and metabolites. A large number of methods have been proposed in the literature for this task. The vast majority of these methods use as input expression levels of the biomolecules under study together with their membership in pathways of interest. The latest generation of pathway enrichment methods also leverages information on the topology of the underlying pathways, which as evidence from their evaluation reveals, lead to improved sensitivity and specificity. Nevertheless, a systematic empirical comparison of such methods is still lacking, making selection of the most suitable method for a specific experimental setting challenging. This comparative study of nine network-based methods for pathway enrichment analysis aims to provide a systematic evaluation of their performance based on three real data sets with different number of features (genes/metabolites) and number of samples.

**Results:** The findings highlight both methodological and empirical differences across the nine methods. In particular, certain methods assess pathway enrichment due to differences both across expression levels and in the strength of the interconnectedness of the members of the pathway, while others only leverage differential expression levels. In the more challenging setting involving a metabolomics data set, the results show that methods that utilize both pieces of information (with NetGSA being a prototypical one) exhibit superior statistical power in detecting pathway enrichment.

**Conclusion:** The analysis reveals that a number of methods perform equally well when testing large size pathways, which is the case with genomic data. On the other hand, NetGSA that takes into consideration both differential expression of the biomolecules in the pathway, as well as changes in the topology exhibits a superior performance when testing small size pathways, which is usually the case for metabolomics data.

## Background

Pathway enrichment has become a standard tool in the analytic pipeline for Omics data, since it reduces the complexity and provides a systems view of the biological question under investigation [1, 2, 3, 4, 5]. Dozens of methods have been proposed in the literature, ranging in modeling sophistication and effectiveness [6, 7, 8, 9, 10, 11, 12, 13, 14, 15, 16, 17, 18, 19]. A number of papers have provided comprehensive reviews of available methods [20, 21, 22] capturing the evolving technical landscape, as well as the range of data types and applications (genes, proteins, etc.). In [20], existing methods have been classified into three generations, the first two corresponding respectively to over-representation analysis (ORA) and functional class scoring (FCS) methods. FCS methods were motivated by the fact that there may be a coordinated activity in functionally related sets of genes, even though each one of them may not be deemed significantly differential by over-representation analysis. The current review focuses on the third generation of pathway analysis methods, namely, topology-based pathway enrichment analysis methods, which utilize information about the interconnections of genes (or other biomolecules) within the pathways, and offer improved performance over conventional second generation methods [6, 7].

Despite the plethora of available methods, there has been a scarcity of systematic comparisons of their performance in controlled settings based on synthetic or real data sets. In [23] and [24], several pathway analysis methods were compared in case studies by assessing the consistency of selected significant pathways. The results confirm that nominated enriched pathways can differ widely across methods, which coupled with absence of a ground truth, makes it difficult to offer guidance to practitioners. In [25], pathway enrichment methods that do not use topology information were compared to topology-based ones. The conclusion was that topology-based methods exhibit superior performance when the pathways under study do not overlap, but not otherwise. However, the data sets examined correspond to small scale studies with no more than 50 samples in total. Moreover, existing comparative studies have primarily focused on methods developed for gene expression data, which may not be suitable for studies involving metabolomics or lipidomics data, where quantitation of all biomolecules in the pathways under study may be incomplete.

Pathway enrichment methods aim to compare the ‘activity’ of pathways of interest across two or more biological conditions or groups of specimens (patients, cell lines, etc.). At the technical level, an important feature is the nature of the statistical null hypothesis being tested. Most methods can be categorized to those testing (I) *self-contained* and (II) *competitive* null hypotheses [26]. A competitive null hypothesis compares the activity of each pathway with other biomolecules/pathways. In contrast, a self-contained null hypothesis compares the activity of each pathway across the biological conditions (e.g. normal vs. disease samples), without comparing it to the other biomolecules/pathways. This difference in the objective results in differences in procedures used for evaluating self-contained and competitive hypotheses (including, e.g., permutation strategies), as well as interpretations of results. In the discussion of methodological issues concerning analysis of sets of biomolecules in [26], the authors argued against the competitive null hypothesis, since tests based on it consider biomolecules as the sampling units which are clearly not independent.

In the current comparative study, we examine nine popular topology-based pathway analysis methods that investigate different null hypotheses. Methods considered in this review have good user interface in R, and include Pathway-Express [8], SPIA [9], NetGSA [10, 11], topologyGSA [12], DEGraph [13], CAMERA [14], CePa [15], PRS [16] and PathNet [17]. Given the importance of availability of open-source software for conducting simulation experiments, popular approaches with only web-based interfaces, such as Ingenuity Pathway Analysis [27], are not included in our comparison. We assess the performance of the above nine methods by performing an extensive numerical analysis using *in silico* experiments that offer advantages over simulation experiments conducted on purely synthetic data [25] or evaluation based on publicly available data [23, 24]. On the one hand, unlike commonly used simulation experiments, our experiments maintain the complexity of real gene/metabolite expression data sets by generating simulated signals from three data sets containing gene and metabolite expressions. On the other hand, unlike comparisons based on publicly available data, active genes/metabolites and pathways are well-controlled in our experiments, allowing us to assess false positive rates and statistical power of the nine pathway analysis methods. Moreover, our review differs from previous ones in that we base our comparisons both on gene expression data and on metabolomic data. The two gene expression data sets studied contain at least 100 samples per condition/group, which enable us to include more genes and pathways of potential relevance in the enrichment analysis. The metabolomic data set provides a different perspective as enrichment analysis of metabolomics data presents additional challenges due to the smaller size of biochemical pathways, their high degrees of overlap, as well as their incomplete coverage by many mass spectrometry acquired data sets.

## Experimental Design

To systematically evaluate the type I error and power of the nine pathway enrichment methods, we consider a variety of settings using three different data sets, two from cancer genomics and one based on metabolomics.

### Pathway dysregulation

We analyzed primary metabolic pathways mapped from KEGG in the metabolomic study, and KEGG pathways that describe signaling and biochemical functions and their interactions in the cancer genomic studies (see Additional File 1 for the complete lists of pathways analyzed). To obtain synthetic, but realistic omics data, a subset *q* < *K* out of a total *K* pathways were first randomly selected as ‘dysregulated’. Next, a pre-specified proportion of genes/metabolites within each dysregulated pathway was chosen to be altered. This proportion, referred to as *detection call* (DC), was set to be 20% for the metabolomic study and 10% for the cancer genomic studies. Due to the smaller pathway sizes and limited number of known interactions within each pathway, affected metabolites were selected randomly from each dysregulated pathway. Since genetic pathways are typically larger in size and have well documented interactions in the `graphite` package, we considered three mechanisms to select affected genes within each dysregulated pathway following the practice in [25].

Note that due to the overlap amongst pathways, a 10% DC threshold may lead to some pathways with more than 10% affected genes/metabolites and more than *q* pathways with at least one affected gene/metabolite.

#### Betweenness

The betweenness of a gene quantifies how often the gene appears on the shortest path between two other genes and effectively measures how important a gene is in the pathway. To select affected genes under this dysregulation design, pathway members are first ranked by their degrees of betweenness. Affected genes are set to be those whose betweenness is above a certain threshold, which is chosen so that a 10% DC is reached.

#### Community

Biological networks tend to contain modules (communities) that are densely connected internally and loosely connected in between. For a pre-specified pathway with a given topology, we find the communities within the pathway using a community detection algorithm, e.g., using the function `cluster_edge_betweenness` from the igraph R package [28]. We then search for a community that approximately represents the 10% DC level.

#### Neighborhood

Under the neighborhood dysregulation design, members within a certain shortest path distance of a randomly chosen gene are used to define the affected genes. The distance parameter is optimized such that a 10% DC is reached after looping over all members within the pathway.

### Data generating model

The steps for generating simulated data are illustrated in Figure 1 and described below. We start with the original log-transformed expression data from *p* genes/metabolites and *n* samples (left heat map in Figure <u>1</u>). Each sample is assumed to follow a distribution *f* with mean *µ* and covariance Σ. More specifically, let

**Figure 1.**
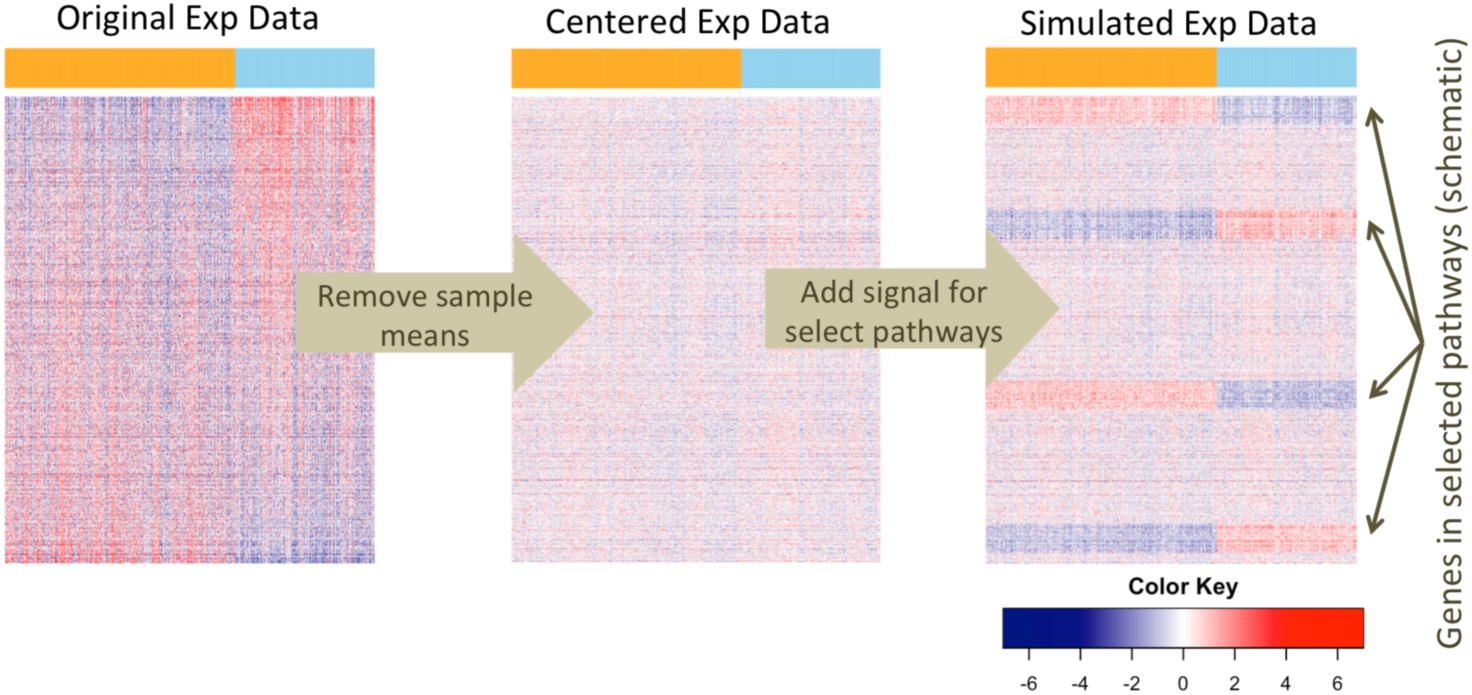
Schematics of the simulation design. In each study, real expression data is used to carry out the simulations. From left to right, the original expression data is first standardized such that each gene/metabolite has mean zero and unit variance. Varying mean signals are then added to genes/metabolites in the selected pathways in each of the simulation replicates. The top bar indicates the sample labels, which are either the original case/control status or a random permutation of the original case/control status.

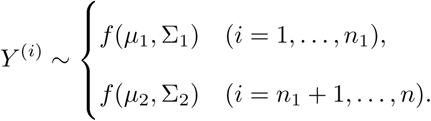

Here we do not assume *f* to be the multivariate normal distribution not only because the distribution of real data is far more complex but because we aim for an agnostic mechanism for data generation that does not favor any method. By assuming a general distribution *f*, we can also assess whether the compared methods are sensitive to normality assumptions.

The data were standardized so that each gene/metabolite has mean zero and unit variance. This corresponds to the middle heat map in Figure 1 and the model

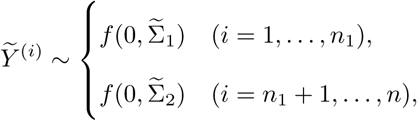

where 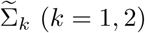 is the correlation matrix.

We considered both settings with and without sample permutation.

I. Sample labels are fixed to be the original case/control status. We added mean signals varying from 0.1 to 0.5 to affected genes/metabolites selected according to different pathway dysregulation designs. This corresponds to the right heat map in Figure 1 and the model

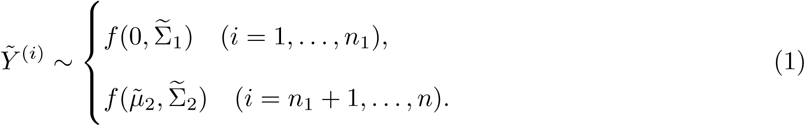
II. We permuted the sample labels first and added varying mean signals to the same set of affected genes/metabolites as in (I). This corresponds to the model

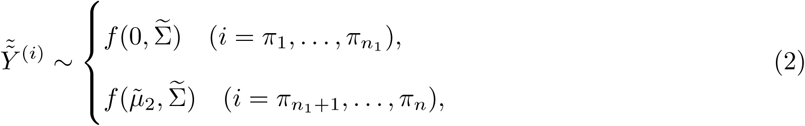

where 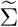 is the common covariance matrix for both populations, and *π* is a random permutation of the sample index 1,…, *n*.

The main difference between models (1) and (2) is whether sample labels are permuted. The intuition is that the permutation version nullifies the difference in covariances, thereby creating a situation where the assumption of equal covariances is satisfied. The permutation version can benefit certain methods, such as DEGraph, but is not ideal for methods such as CAMERA that exploit difference in correlations. Theoretical justifications for how permutation works are available in [29] and [30].

The above data generating mechanism relaxes the distributional assumptions and allows us to evaluate different methods with respect to a ground truth. This is in contrast to most existing reviews in the literature based on either publicly available data where the ground truth is unknown [23, 24], or purely synthetic data from parametric distributions [25]. In practice, because there is only one deterministic data matrix, for both (I) and (II) we added independently and identically distributed Gaussian noise to each entry in the data matrix to introduce randomness.

The complete set of test designs is summarized in Additional File 1.

## Results

We present the results of our comparative study by assessing the performance of the nine methods in terms of their type I errors, followed by their statistical powers. A type I error, also called a false positive, occurs when a true null hypothesis is rejected, whereas the power quantifies the probability of a test correctly rejecting the null when the alternative hypothesis is true. Of course, power comparison is only meaningful if the tests all have valid type I error control. Due to the large size of the two gene expression data sets, the type I errors and powers were calculated as the proportion of null hypotheses rejected among 200 simulation replications, but they were evaluated over 1000 replications in the metabolomic data example.

Several methods require *p*-value thresholding. To this end, we used univariate two-sample *t*-test assuming unequal variances to calculate the *p*-value for each gene/metabolite. These *p*-values were corrected for multiple comparisons using the Benjamini & Hochberg procedure for controlling false discovery rate (FDR) [31]. The FDR cutoff of 0.05 was used to identify DE genes/metabolites.

In our analysis, multiple method-specific difficulties made it impossible to compare all available pathways. First, in the two gene expression data examples, SPIA and Pathway-Express returned *p*-values primarily for signaling pathways, which is expected given their testing procedures. Second, topologyGSA only works for pathway whose topology is a DAG and whose size is smaller than the minimum number of samples in the two conditions/groups. As a result, our comparisons in the two cancer genomic applications only include selected pathways that were analyzed by all methods. Lastly, the network in the metabolomics data example is not directed acyclic and the pathways are smaller biochemical pathways that often have incomplete coverage and high degrees of overlap. Thus we excluded topologyGSA, Pathway-Express, SPIA and PRS in the metabolomics data example.

### Ranking empirical powers

In power comparisons, we summarize the relative performance of all nine methods based on the ranking of each pathway’s empirical power. Specifically, for each pathway, we rank the empirical powers of all methods at each level of mean change from low (indicating higher power) to high (indicating lower power). Methods that produce ‘NA’ (e.g. ORA methods when there are too few DE genes) are ranked the highest. For each pathway, the geometric mean of all rankings across different mean changes is taken as the final measure of relative performance. Spreadsheets with disaggregated empirical powers for each pathway, across all the experimental settings, are provided in [32].

### Analysis of gene expression data

There are 11 KEGG pathways (primarily on signaling) whose topologies satisfy the input requirement needed for topologyGSA and Pathway-Express. We thus focused on type I error and power comparisons on this subset of pathways under the betweenness dysregulation design. Details on the pathways and more comparisons are available in Additional File 1.

The scatter plot in Figure 2 shows the type I error for each of the 11 KEGG pathways using the TCGA breast cancer data [33]. Each point indicates the type I error rate for one pathway. Overlapping points were re-positioned by adding ±1 to the x-axis and ±0.01 to the y-axis, which explains why some values are less than zero—these should be understood as being close to zero. Because the number of DE genes under the self-contained null is zero even with liberal FDR cutoffs, ORA-type methods such as SPIA, CePa, PRS and PE.Cut (Pathway-Express with *p*-value cutoff) can not assess the pathway significance. These methods have thus been excluded from type I error comparison. On the other hand, PE.noCut does not require *p*-value thresholding and is therefore included in Figure 2. Across the 11 pathways, all methods control the type I error rate at 0.05 significance level. It is worth noting that most of the type I error rates from PE.noCut and PathNet are close to the nominal level 0.05, whereas all other methods seem to have conservative type I error rates.

**Figure 2.**
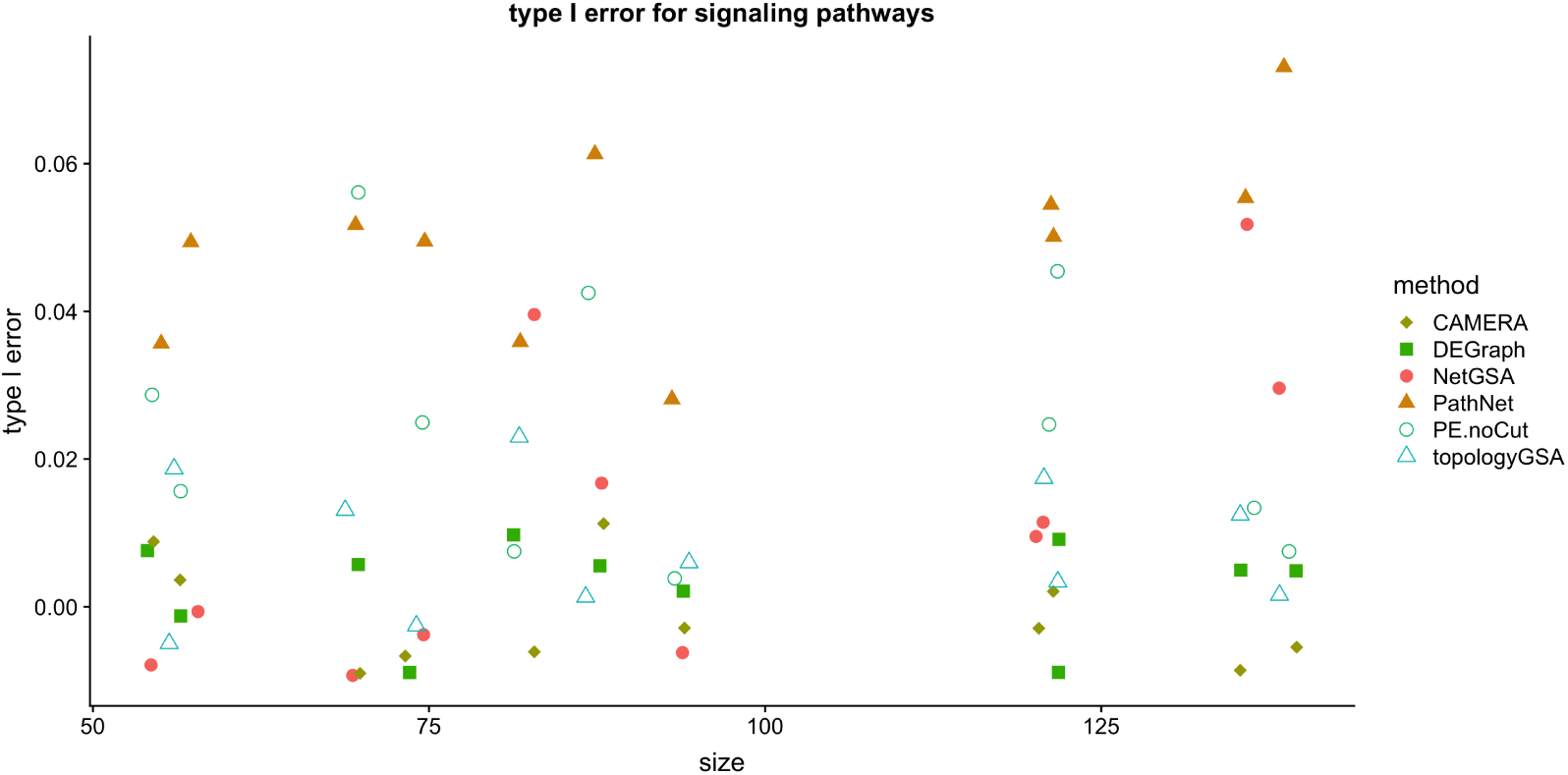
Type I errors for the 11 KEGG (primarily signaling) pathways in the TCGA breast cancer study [33]. The x-axis shows the pathway size and the y-axis indicates the type I error. Overlapping points were re-positioned by adding ±1 to the x-axis and ±0.01 to the y-axis. Thus the negative values should be understood as being very close to zero. All methods control type I errors.

Figure 3 presents the relative performance of different methods in terms of average ranking of empirical powers for the 11 pathways, both with and without sample label permutation. Lower rankings indicate better performance. Again we re-positioned overlapping points by adding ±1 to the x-axis and ±0.1 to the y-axis for visualization purpose, which explains why some rankings in Fig 3A are below 1. Among all methods, PathNet, CAMERA and PE.noCut seem to perform the best when using the original sample labels (Fig 3A). NetGSA and DEGraph have similar performance. On the other hand, when sample labels are permuted, DEGraph, PE.noCut and topologyGSA have the best performance (Figure 3B). This is not surprising because the permutation version nullifies the difference in covariances asymptotically, thereby creating a situation where the assumption of equal covariances and shared graph structure is approximately satisfied. DEGraph and topologyGSA thus performed better relative to the others, whereas the performance of CAMERA deteriorated, especially for large pathways, because it cannot leverage the difference in correlations. Sample permutation does not affect PE.noCut because its significance is evaluated based on gene permutation. All ORA-type methods (SPIA, PE.Cut, CePa and PRS) have relatively higher ranking and hence poorer performance because they only work when the magnitude of mean changes between conditions is large.

**Figure 3.**
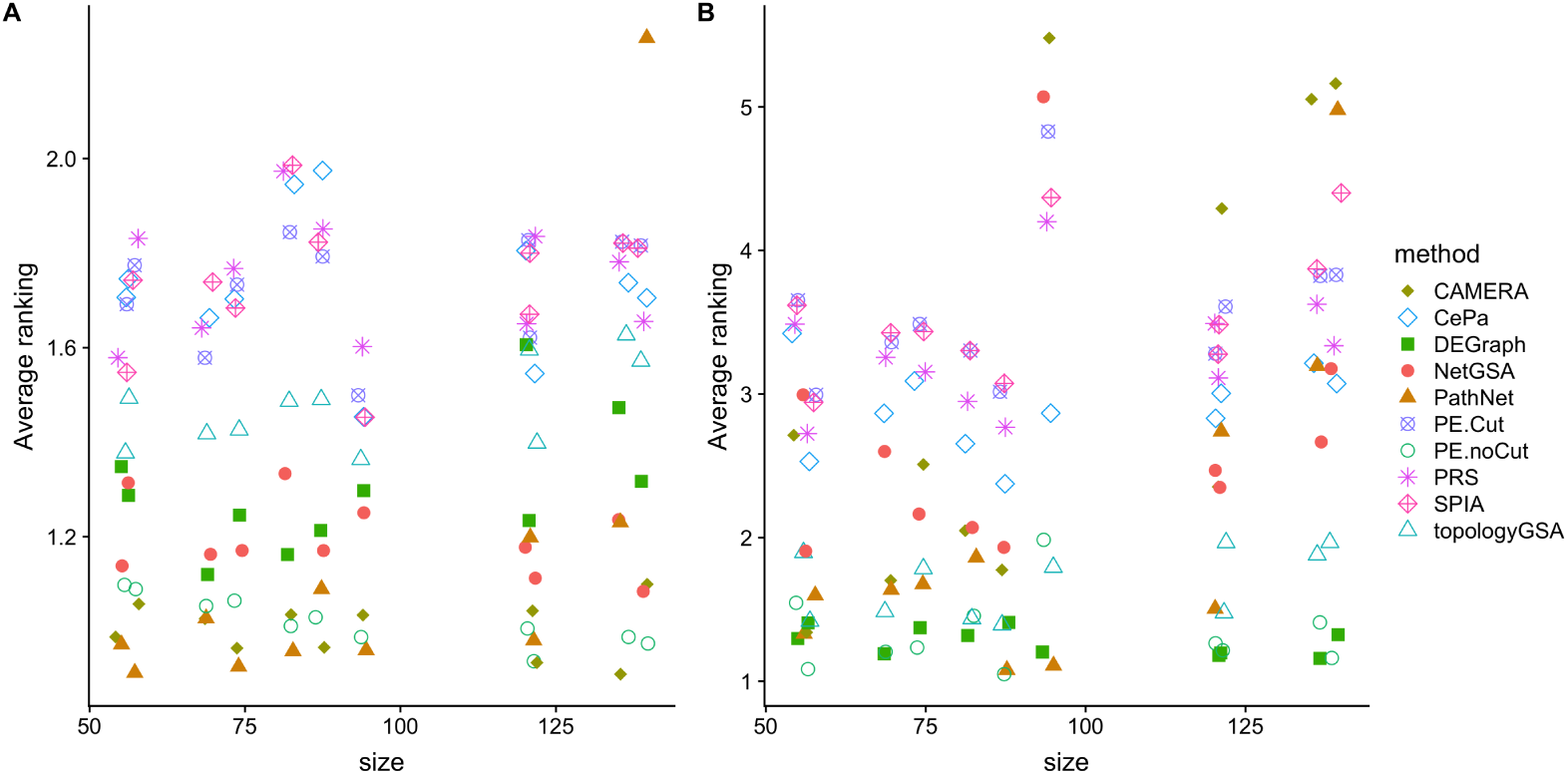
Average ranking of empirical powers on the 11 KEGG (primarily signaling) pathways using sample labels from the original study (A) and shuffled sample labels (B) based on the betweenness dysregulation design for the TCGA breast cancer study [33]. The x-axis shows the pathway size, and the y-axis indicates the average ranking of empirical power over different mean changes. Lower ranking indicates better performance. Overlapping points were re-positioned by adding ±1 to the x-axis and ±0.1 to the y-axis. PathNet, CAMERA and PE.noCut perform the best when using the original sample labels (A), whereas DEGraph, PE.noCut and topologyGSA yield the best performance with permuted sample labels (B).

Similar figures for other dysregulation designs and other pathways, including those for the prostate cancer study, are available in Additional File 1. It is worth noting that PathNet shows inflated type I error rates on several KEGG metabolic pathways (Figures S1 and S4), which could be due to the inaccurate coverage in pathway topology in `graphite`. Methods such as topologyGSA, SPIA, Pathway-Express and PRS are not applicable to those metabolic pathways due to constraints in their topology. Among the methods compared, DEGraph has the best overall performance.

### Analysis of metabolomics data

The findings in the metabolomics data example differ qualitatively from those in genomic examples. This difference can be attributed to the relatively small number of edges in the metabolic network and small pathway sizes, which are due to the incomplete coverage of metabolomic assays. Note because SPIA, Pathway-Express and PRS were proposed specifically for genetic pathways, and the metabolic network is not directed acyclic, we only compared NetGSA, DEGraph, CAMERA, CePa and PathNet in this example. In addition, CePa requires the presence of DE metabolites, which only appear when the mean change is greater than 0.5. The mean signal in this example was thus allowed to vary from 0.1 to 1.0. To run all 5 methods, we filtered out 33 of the 65 KEGG pathways whose topologies are too sparse—fewer than two edges—and focused on type I error and power comparisons on the remaining 32 pathways.

Figure 4 shows that all four methods control type I errors, although NetGSA and CAMERA exhibit slightly inflated type I errors for some pathways, which is likely due to the small sample sizes available. This will be less of an issue for NetGSA if more samples are available for network estimation; such samples can be obtained from other related studies. The conservative type I errors of DEGraph is again due to its assumption that the networks for the two conditions are the same. Since there are very few pathway interactions available in the metabolic network, PathNet also exhibits conservative type I errors.

**Figure 4.**
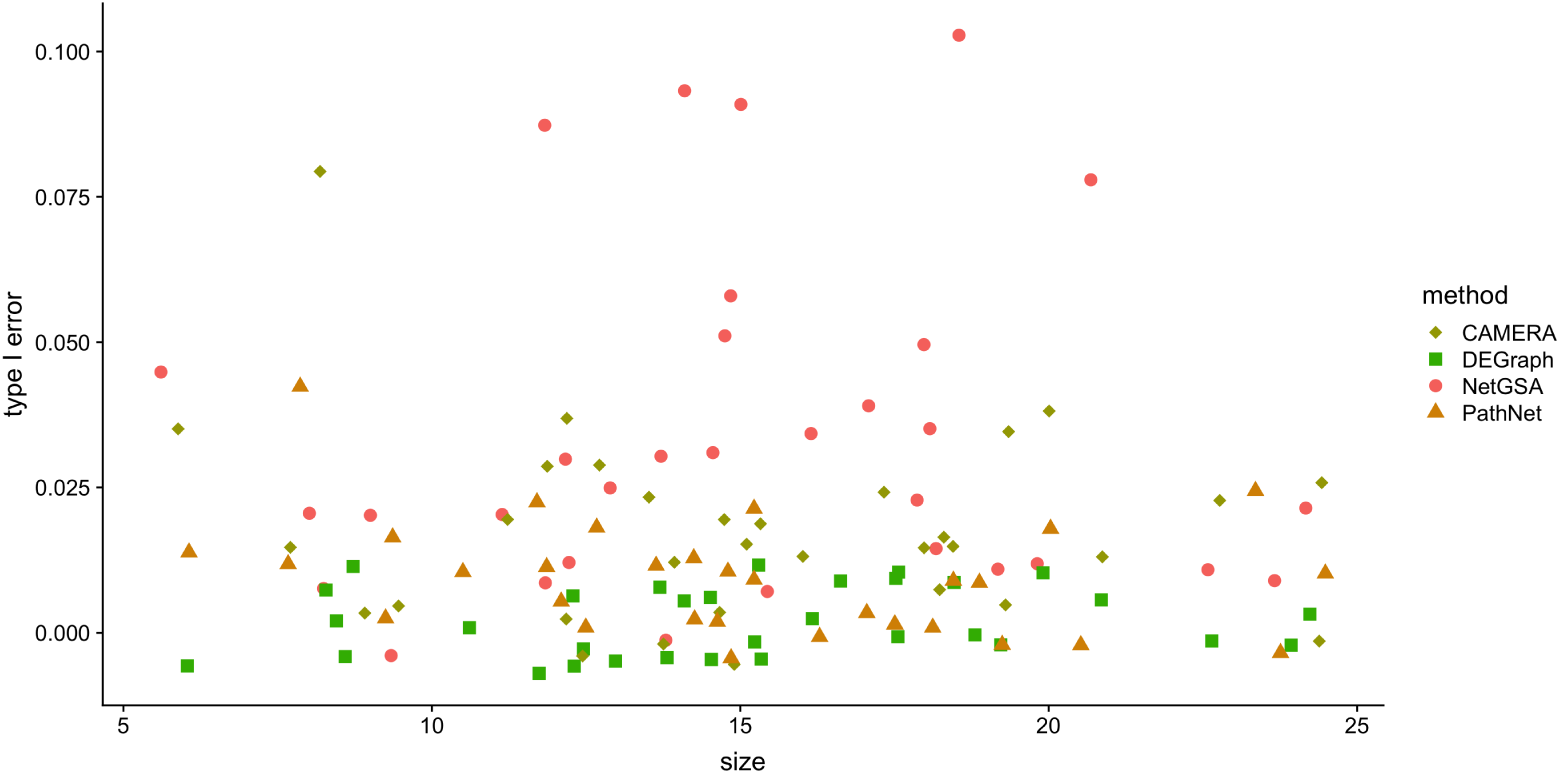
Type I errors for KEGG metabolic pathways in the metabolomics data [39]. The x-axis shows the pathway size and the y-axis indicates the type I error. Overlapping points were re-positioned by adding ±1 to the x-axis and ±0.01 to the y-axis. CAMERA, DEGraph and PathNet have controlled type I errors, but NetGSA’s type I errors are slightly inflated for several pathways.

Figure 5 compares the relative performance of different methods in ranking of empirical powers out of 1000 replications. It is evident that DEGraph and NetGSA perform the best (lowest rankings), regardless of whether the sample labels are shuffled. DEGraph performs slightly better than NetGSA, especially under permuted sample labels (Figure 5B), because the Hotelling’s T-squared test works well for small pathways. In comparison, the linear mixed effects model underlying NetGSA considers an additional random effects, resulting in slightly lower power when samples are permuted. The other methods CAMERA, CePa and PathNet have comparable performances both with and without permuting sample labels. The poor performance of PathNet and CePa is due to the sparse metabolic network, whereas CAMERA does not work well because there is high overlap among metabolic pathways (Figure S7 in Additional File 1).

**Figure 5.**
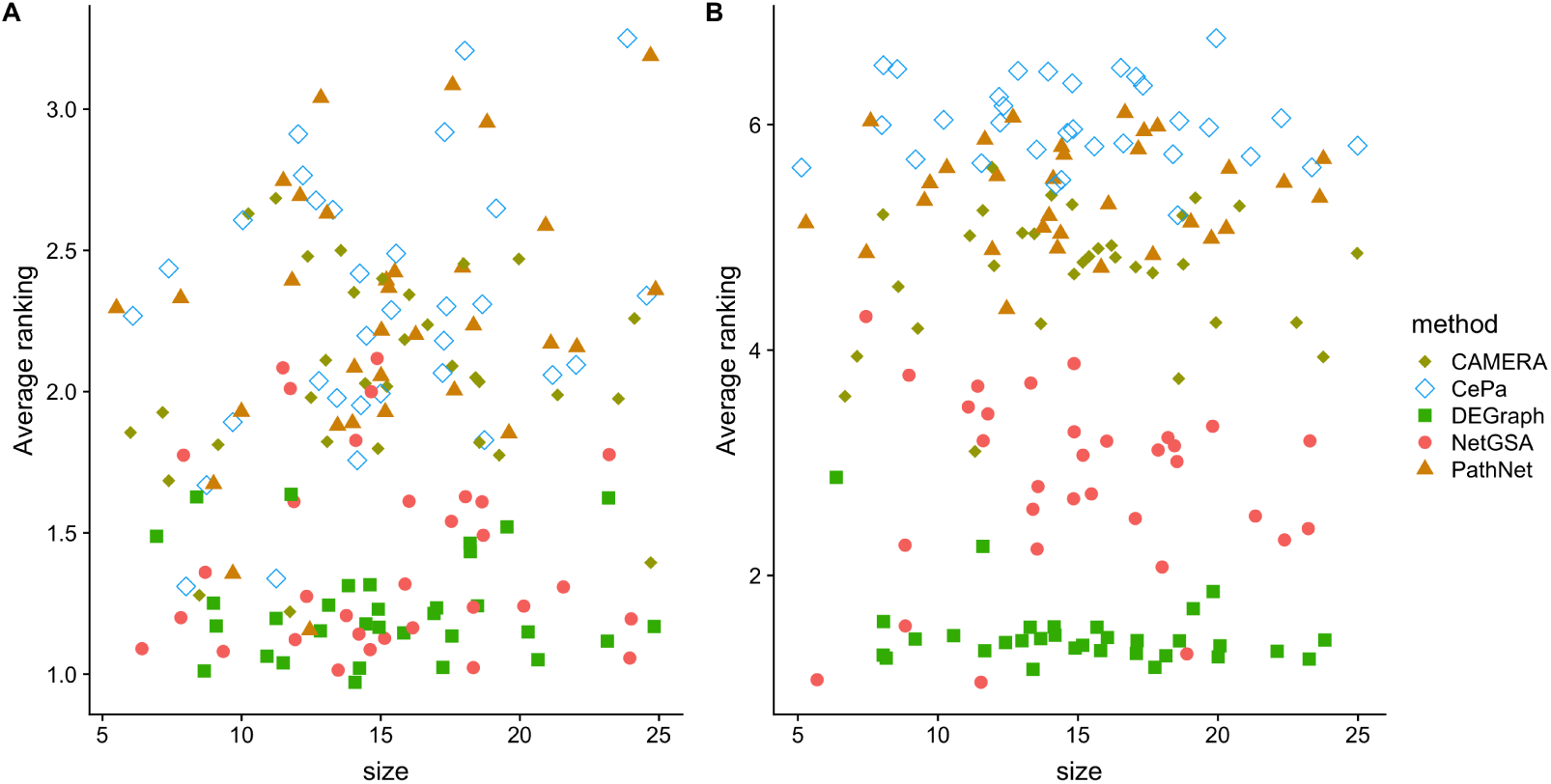
Average ranking of empirical powers using sample labels from the original study (A) and shuffled sample labels (B) for the metabolomics data [39]. The x-axis shows the pathway size, and the y-axis indicates the average ranking of empirical power over different mean changes. Lower ranking indicates better performance. Overlapping points were re-positioned by adding ±1 to the x-axis and ±0.1 to the y-axis. DEGraph and NetGSA perform the best regardless of sample permutation.

## Discussion

The methods considered in this study exhibit differences in terms of the network information that they incorporate. For example, CePa, PathNet, Pathway-Express, PRS and SPIA only account for the pathway topology. CAMERA does not directly take into account the pathway topology, but estimates the correlations among the biomolecules from data. In contrast, NetGSA, DEGraph and topologyGSA assume the underlying networks come from a known class of graphs, whose parameters are inferred from data. NetGSA can also incorporate existing network information from user-provided sources for improved power. The downside with the more flexible version of NetGSA that accounts for differences in networks is that the type I errors may be slightly inflated, if too few samples are available for the estimation of the network parameters, corresponding to network edges. This was observed in the metabolomics data example. This issue can be addressed by aggregating data from multiple related studies for more accurate network estimation.

Taking a broader view point, NetGSA is capable of assessing pathway enrichment due to changes both in the mean levels of the biomolecules, as well as their connectivity; it can thus be more suitable for studies involving comparisons across different disease states, where a possible strong dysregulation of the interactome can occur. It can also seamlessly accommodate more than two conditions [34], as well as multiple types of Omics data for integrated analysis of pathway enrichment [35]. Although fairly robust in the examples shown above, a disadvantage of DEGraph is that it is not particularly suitable in settings where one expects a strong dysregulation of the underlying interactome, since it assumes the underlying networks in different conditions have the same structure. Consider, e.g., the two networks in Figure S8 of Additional File 1. The proportion of nodes in pathways 8, 4 and 7 that have nonzero mean changes is 0.2, 0.4 and 0.6, respectively. In contrast, pathways 3, 5, and 2 have not only the corresponding level of mean changes, but also complete rewiring of the pathway topology. In this simulated setting, NetGSA is able to identify pathways 5, 3 and 7 as the most significantly enriched pathways (with empirical power at least 0.9), followed by pathway 2 (0.89). DEGraph identifies pathways 2, 3 and 5 as being enriched, but misses pathway 7 (0.62).

Importantly, existing methods such as CAMERA, CePa, DEGraph, PathNet, Pathway-Express, PRS, SPIA and topologyGSA often analyze one pathway at a time while ignoring the fact that pathways overlap with each other. This practice is common because it is conceptually and computationally convenient, yet it may lead to undesirable consequences as the interactions between genes/metabolites that show up in multiple pathways may be different when analyzing these pathways separately. Separate analysis is particularly problematic for metabolomic studies where the pathways are considerably smaller in size and exhibit a high degree of overlap (Figure S7). In contrast to these methods, NetGSA can combine all metabolites by inferring a joint network, and is thus more powerful for detecting enriched pathways. However, simultaneous analysis of all pathways using NetGSA could become computationally expensive, if the study contains a very large number of biomolecules.

## Conclusions

Significant progress has been made in developing topology-based methods for pathway enrichment analysis. In this study, we undertook a systematic comparison of nine popular such methods using three data sets from gene expression and metabolomics profiling. Compared to existing reviews [23, 24, 25, 36], our comparison leverages the large sample sizes in the two cancer genomic studies, and, in particular, offers important insights for how the nine competitors perform in metabolomics studies, where the focus is on smaller biochemical pathways. Results in Additional File 1 (Figures S2, S3, S5, S6) suggest similar findings as those observed in Figure 3, and confirm the overall robust performance of DEGraph. In general, when the methods examined are used for pathway enrichment purposes in studies based on genomic data and focusing on large signaling pathways, most exhibit a satisfactory overall performance, with DEGraph being the most robust, followed by PathNet, Pathway-Express without *p*-value cutoff (PE.noCut) and topologyGSA. In comparison, ORA-type methods (SPIA, PE.Cut, CePa and PRS) require the presence of DE genes and perform well only in specific settings. Our experience with Pathway-Express is that PE.noCut (without *p*-value cutoff) seems to dominate PE.Cut (with *p*-value cutoff). Additionally, due to inaccurate coverage and special features of topology information on smaller metabolic pathways, only DEGraph shows robust performance (Figures S3, S6). On the other hand, in studies involving metabolomics data and pathway enrichment of relatively small biochemical pathways, NetGSA and DEGraph clearly outperform all competitors.

## Methods

In this section, we describe the three data sets used in our comparative study and then provide an overview of the nine topology-based pathway enrichment methods analyzed.

### Data sets

Our first comparison considers a breast cancer gene expression study from The Cancer Genome Atlas [33, TCGA]. We focused on 114 signaling and metabolic pathways from the Kyoto Encyclopedia of Genes and Genomes [37, KEGG], and expression data from 2784 genes that have matched Entrez IDs. The data set consists of 520 samples in total, 117 estrogen-receptor-negative (ER-) and 403 estrogen-receptor-positive (ER+).

Our second comparison is based on a TCGA gene expression data on prostate cancer from [38]. The data contained Affymetrix probe IDs, which were first mapped to gene Entrez IDs. When multiple probes mapped to a single gene (i.e., to the same gene Entrez ID), their mean profile was used to avoid duplicated gene IDs. To reduce the dimensionality, genes in 112 KEGG signaling and metabolic pathways were considered for the final analysis. The final data set contains expression levels of 2952 genes across 264 case and 160 control subjects.

The third and final data set comes from a metabolomics study on non-obese diabetic mice, where the metabolic profiles of 41 non-diabetic and 30 diabetic animals of 100 named metabolites were collected, with the goal of identifying metabolic signatures of Type I diabetes progression [39].

### Pathway enrichment methods

#### Pathway-Express

The Pathway-Express method [8] for analyzing signaling pathways is implemented in the ROntoTools Bioconductor package [40]. Its null hypothesis is that the list of DE genes on a given pathway is completely random; this hypothesis is tested by calculating an impact factor for each pathway *G* defined as

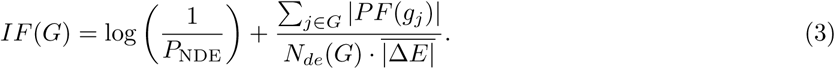

Here, *P*_NDE_ in the first term on the right hand side of (3) evaluates the significance of the pathway *G* as measured by an over-representation analysis, whereas the second term incorporates the DE genes in the data and the interactions among genes inside the pathway. The perturbation factor (PF) for each gene is defined as

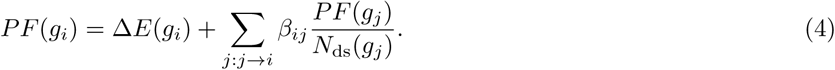

In Equation (4), Δ*E*(*g*_*i*_) is the signed normalized expression change of biomolecule *g*_*i*_. The denominator *N*_ds_(*g*_*j*_) represents the number of downstream biomolecules of *g*_*j*_ and the sum is only over biomolecules *g*_*j*_ directly upstream of *g*_*i*_. The interactions between *g*_*i*_ and *g*_*j*_ are encoded in the coefficients *β*_*ij*_, e.g. +1 for activation, −1 for inhibition. Therefore, the second term on the right hand side of (3) can be thought of as the total perturbation factor normalized by (i) the number of DE genes in pathway *G*, which is *N*_*de*(*G*)_, and (ii) the mean absolute fold changes among all DE genes in the data, which is 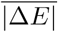. Normalization with respect to 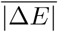 is to account for the potential differences in estimating the fold changes among various technologies. The random quantity *IF* (*G*) is shown to have a Γ(2, 1) distribution, such that for any realization of *IF* (*G*) = *f* the significance of a pathway can be calculated as

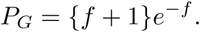

Pathway-Express is implemented in the R package ROntoTools. The latest version of this package (2.10.0) also permits a cutoff-free version which eliminates the need to select DE genes [41]. In this version, Pathway-Express does not calculate the ORA significance, but only reports the significance from pathway perturbation.

#### SPIA

Signaling pathway impact analysis [9, SPIA] tests the same null hypothesis as Pathway-Express; it combines one evidence based on *P*_NDE_ with a second evidence, *P*_PB_, that quantifies the amount of perturbation in each pathway. To calculate the second type of evidence, the total net perturbation accumulation for a given pathway *G* is defined as

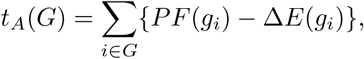

where the perturbation factor *PF* (*g*_*i*_) is defined in (4). A bootstrap approach is used to obtain the perturbation *p*-value *P*_PB_, which is the probability of observing a total accumulated perturbation of a pathway more extreme than *t*_*A*_(*G*) just by chance.

The overall significance of pathway *G* is calculated as

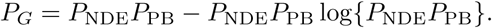

It is worth noting that Equation (4) imposes an implicit constraint on the pathway topology, in that a pathway with a singular matrix *I* −*B* for *B*_*ij*_ = *β*_*ij*_*/N*_ds_(*g*_*j*_) cannot be analyzed using Pathway-Express and SPIA. In addition, both Pathway-Express and SPIA require the presence of DE genes to define the impact of pathways. Hence pathways that do not have any DE genes will not be analyzed. This phenomenon was observed in [9, 36] and in our comparisons, where Pathway-Express and SPIA often only return the significance of half of all pathways considered.

#### NetGSA

The NetGSA method [10, 11] employs as input directed and/or undirected networks that define pathway interconnectedness. If the network information is incomplete, it uses a probabilistic graphical model to complete the pathway topology based on the available data, while using the existing topology information as constraints. As a result, not only can NetGSA estimate novel interactions, but also validate existing network information. Next, we present the statistical model used in NetGSA assuming the underlying networks are undirected.

Given the adjacency matrix *A* of the network, NetGSA defines the propagated effect of genes on each other through the *influence matrix* Λ, defined as ΛΛ^*′*^ = (*I*−*A*)^−1^ with *I* denoting the identity matrix. It then decomposes the measurements in the *i*th sample, *Y* ^(*i*)^, into signal *X*^(*i*)^ and noise ***ε***^(*i*)^. The multivariate signal’s interactions are captured through a Gaussian Markov random field encoded by *A*. Formally, let ***γ***^(*i*)^ be the baseline expression levels of all biomolecules and ***µ*** be their mean expression levels. NetGSA decomposes *X*^(*i*)^ = Λ***γ***^(*i*)^ so that the signal for the *j*th gene, 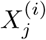, combines both its baseline activity 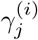 and those propagated from its neighbors. Given data from *K* conditions, NetGSA allows for the *K* networks to be different and for each condition *k* considers a linear mixed effects model

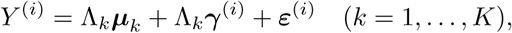

where the random effects 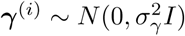 are independent from noise ***ε***^(*i*)^.

Let ***β*** be the concatenated vector of the baseline means ***µ***_1_, …, ***µ***_*K*_. To test for enrichment of any pathway *G*, NetGSA uses a Wald test statistic,

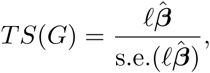

for *K* = 2 or an F statistic for *K* ≥ 3 to test the null *𝓁****β*** = 0; here, 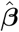 denotes the estimate of ***β*** based on the data, s.e. 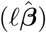 represents the standard error of 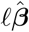, and *𝓁* is a contrast vector optimally defined to allow for simultaneous testing of differences in the mean structure across the *K* conditions, as well as differences in interaction networks.

NetGSA can be computationally slow in the presence of a large number of biomolecules, which is the case for the two studies involving gene expression data. In those instances, enrichment analysis is carried out separately for each pathway. On the other hand, since the metabolomic data set contains only 100 metabolites, enrichment analysis of all pathways is performed simultaneously.

#### topologyGSA

The pathway topology information in the topologyGSA method [12] is first converted into a directed acyclic graph (DAG) and then to its moral graph, which represents its corresponding Markov equivalence class [42]. Let the data be organized such that the first *n*_1_ columns correspond to samples from condition 1, and the last *n* − *n*_1_ columns from condition 2. Given the graph structure, topologyGSA models each sample *Y* ^(*i*)^ using a probabilistic graphical model approach:

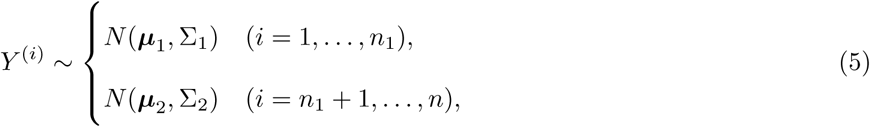

where ***µ***_*K*_ is the mean expression level for condition *k*, and Σ_*K*_ is the corresponding covariance. Note Σ_1_ and Σ_2_ are constrained to have the same structure as specified *a priori*. topologyGSA first tests the hypothesis of equal variances Σ_1_ = Σ_2_ using a likelihood ratio test. Depending on the conclusion from testing the equality of variances, the test of differential expression ***µ***_1_ = ***µ***_2_ is performed through a multivariate analysis of variance (MANOVA) [43] if Σ_1_ = Σ_2_, or based on the Behrens-Fisher problem [44] if Σ_1_ ≠ Σ_2_. The method is designed specifically for gene expression data.

topologyGSA has several limitations. First, it requires the pathway topology to be organized as a DAG, which may not be possible for, e.g., metabolomics studies. In addition, topologyGSA relies on the likelihood ratio statistic for testing equal covariances, which restricts its use to relatively small pathways. For large pathways with more members than the sample size—which are frequently observed in studies involving gene expression data—topologyGSA can become computationally very inefficient. Lastly, differential network and differential expression are tested separately in topologyGSA, which may also limit the power of the test when the two populations differ in both means and variances.

In our data examples, we implemented both test on differential network using `pathway.var.test` and test on differential expression using `pathway.mean.test` from the `R` package `topologyGSA`. Since naive combination of the two tests (e.g. by taking the minimum *p*-value) resulted in inflated type I errors in our numerical analyses, we used the *p*-value for testing equality of means as representing significance for pathway enrichment.

#### DEGraph

DEGraph was introduced in [13] to conduct a two-sample test of means, while incorporating topology information of the biomolecules. It considers a special case of the model in (5) with Σ_1_ = Σ_2_ and tests the null ***µ***_1_ = ***µ***_2_. The motivation underlying DEGraph is that the classical Hotelling’s *T* ^2^-test [45], which is known to be uniformly most powerful against global mean-shift alternatives for multivariate normal distributions and may behave poorly in high dimensions. When the graph *G* capturing interactions of the biomolecules in the two conditions is known, [13] derived an equivalent expression for Hotelling’s *T* ^2^ statistic in the graph-Fourier space [46]. It further proposed to approximate Hotelling’s *T* ^2^ by filtering out high frequencies of the Fourier coefficients when the dimension is high. The statistic after filtering is shown to yield a test that is more powerful than testing in the original unfiltered space.

Because DEGraph is a test in the graph-Fourier space, it requires knowledge of a connected graph, which is assumed to be the same between the two conditions under consideration. If a pathway consists of more than one connected component, DEGraph will test whether the means are different for each connected subgraph, and correct for multiple comparisons using a permutation procedure. In addition, if the input pathway topology can not be immediately used in constructing a test in the graph-Fourier space, DEGraph offers the functionality of subgraph discovery. In our implementation, we supplied DEGraph with the pathway information from KEGG and let the method decide whether to undertake subgraph discovery.

#### CAMERA

Correlation Adjusted MEan RAnk gene set test [14, CAMERA] is a competitive biomolecule set testing procedure and is available as a function in the limma package [47]. CAMERA assumes that the log-expression value *Y*_*g*_ for biomolecule *g* is linear in the design variables specifying the conditions with coefficients ***α***_*g*_. Enrichment analysis of a given pathway *G* is done by testing the null *𝓁****α***_*g*_ = 0, where *𝓁* is a contrast vector specified by the user. Denote by *z*_*g*_ the biomolecule-level statistic for *g*. Given *m* such statistics *z*_*G*_ = (*z*_1_, …, *z*_*m*_) in pathway *G*, CAMERA tests whether their mean expression 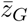 inside a pathway is significantly different from the mean expression of biomolecules outside the pathway. Let *p* be the total number of biomolecules, both inside and outside the pathway, and 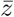 be the mean of all biomolecule-level statistics. CAMERA uses the statistic

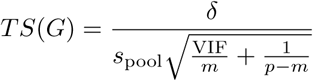

to test the competitive null hypothesis; here, 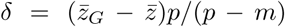 is the adjusted mean difference, *s*_pool_ is the pooled residual standard deviation and 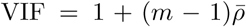 denotes the variance inflation factor. The parameter 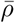 is defined as the average correlation amongst the biomolecule-level statistics inside the pathway and is estimated from the data. Note that CAMERA incorporates the pathway membership information and does not take interconnectedness inside the pathway into account. However, it does account for correlation among biomolecules in the pathway.

#### CePa

CePa is a centrality-based pathway enrichment analysis method [15, 48]. CePa allows multiple centrality measures to capture the topology of a given pathway from different aspects. It also maps genes to pathway nodes and considers the node as the basic pathway unit, which is particularly useful for enrichment analysis of complexes or protein families.

For a given pathway with *m* nodes, the CePa score is defined as

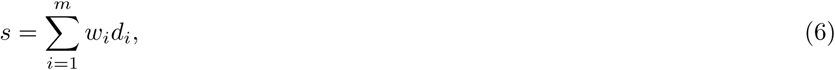

where *d*_*i*_ = 1 if the *i*th node is differentially expressed and *d*_*i*_ = 0 otherwise, and *w*_*i*_ corresponds to the *i*th node’s weight defined based on various centrality measures. A small offset of 0.01 is added to each *w*_*i*_ to ensure positive weights. CePa allows equal weights and weights defined from centrality measures such as node degree, betweenness [49] and the largest reach. The degree centrality measures the number of neighbors a node has, betweenness quantifies how often a node appears on the shortest path between two other nodes, and the largest reach centrality defines how far a node can send or receive information within the pathway.

CePa uses gene permutation to test whether genes inside the pathway are at most as differentially expressed as those outside the pathway given its score. In our numerical results, we took the minimum of the *p*-values obtained using different weights as representing significance for pathway enrichment. The score definition in (6) is a variant of ORA, but can also be extended to incorporate gene-level statistics as in FCS methods [15, 50].

#### PRS

The Pathway Regulation Score [16, PRS] enrichment method was developed in parallel to CePa and the two share some similarities. Specifically, PRS assigns a value *v*_*i*_ and weight *w*_*i*_ to each node *i*, which may contain one or more genes. The node value *v*_*i*_ is 0 if the corresponding gene(s) are not expressed, 1 if they are expressed but not differential, or the maximum fold-change value if one or more genes in node *i* are differential. If *v*_*i*_ *>* 1, node *i* is then assigned a weight *w*_*i*_, which is the number of downstream DE nodes (either directly or via other significant nodes, including the starting node itself). The score for a pathway with *m* nodes is defined as

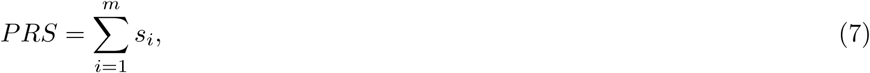

where node score *s*_*i*_ = *w*_*i*_*v*_*i*_ if node value *v*_*i*_ *>* 1 and 0 otherwise. The pathway score *PRS* is then normalized with respect to pathway size by multiplying the proportion of DE genes over the total number of expressed genes.

PRS assesses the significance of each pathway using gene permutation. The raw and permuted scores, calculated respectively from the original data and permuted data, are first standardized with respect to the permuted scores in order to derive the empirical null distribution. The *p*-value of each pathway is determined as the proportion of normalized permuted scores greater than or equal to the normalized raw scores.

#### PathNet

PathNet [17] combines all pathways under consideration into a *pooled* pathway, defined as the union of all pathways. The interactions among genes in the pooled pathway are represented by an adjacency matrix *A*, which is a binary matrix with 1 and 0 indicating the presence and absence of an interaction. Given the network *A*, PathNet calculates the biomolecule-level significance 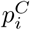 by combining the direct evidence 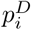 with the indirect one 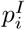 based on Fisher’s method. It then uses a hypergeometric test to evaluate the significance of a given pathway. While the direct evidence 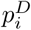 accounts for the differential expression of each biomolecule *g*_*i*_, the indirect evidence 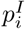 incorporates the effects on *g*_*i*_ from its neighbors. Specifically, PathNet defines the indirect evidence score for *g*_*i*_ as

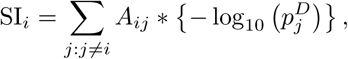

where the sum is over all biomolecules in the pooled pathway. The significance of the indirect evidence 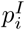 is then determined by testing if the observed SI_*i*_ is greater than expected by chance. Similar to SPIA, PathNet accounts for biomolecule interactions only through the topology information available in the database, e.g., KEGG.

### Methodological considerations

Table 1 provides an overview of all methods in terms of their null hypotheses and input requirements. These methods differ in two main aspects.

**Table 1.**
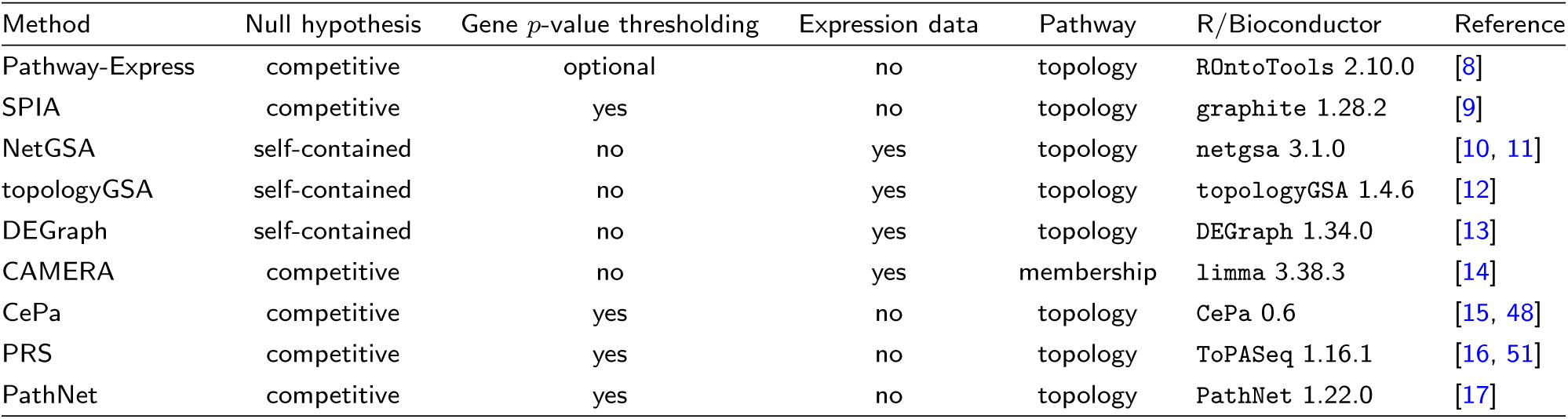
Overview of tested pathway enrichment methods. All methods return the p-values before and/or after correcting for multiple comparisons.

The first distinction is in the type of null hypothesis. CAMERA and PathNet test a competitive null hypothesis of whether the genes in a given pathway are at most as differentially expressed as those outside the pathway. Pathway-Express, SPIA, CePa and PRS test the competitive null by comparing the pathway of interest to a random pathway (while holding the sample labels fixed). In contrast, NetGSA, topologyGSA and DEGraph consider the self-contained null hypothesis by testing a pathway against itself. Although the competitive null hypothesis can have an appealing interpretation, assessing the significance of the competitive null is challenging, as it corresponds to a gene sampling framework which treats genes as independent (see discussion in [26]).

Another major difference among these methods is whether the method takes as input expression data or thresholded gene *p*-values. With the exception of CAMERA, all methods based on testing the competitive null need to determine DE genes based on a pre-specified threshold of corrected *p*-values. While the thresholding may seem intuitive, the resulting enrichment results may be sensitive to the *p*-value cut-off. The emphasis on *p*-value thresholding also implies that these methods will not work in settings where there are too few DE genes, i.e. when gene-wise expression change is too small, as illustrated in the Results section. All self-contained tests directly use expression data and thus avoid making subjective choices about DE genes.

### Network information

In evaluating these methods, we also make the distinction between pathway topology and pathway membership, where the former refers to both pathway membership and the interactions amongst pathway members.

Importantly, the pathway topology information required by different methods can be quite different, which may affect the user experience. CAMERA only uses pathway membership and requires the least effort. The `R` package `graphite` provides functionality to retrieve the list of KEGG pathways, and the resulting topology information can be readily passed to Pathway-Express, SPIA, topologyGSA, DEGraph and PRS (as implemented in `ToPASeq`).

In comparison, NetGSA, CePa and PathNet require additional steps of processing before `graphite` pathways can be analyzed. However, this additional step also implies flexibility in the sense that the user can specify desired network information. For example, NetGSA requires a weighted network, where the weights reflect the interactions between genes/metabolites. This can be either available network connectivity information from a database, or estimated from data based on partial correlations complemented with connectivity information from a database. In the gene expression data examples, we used gene-gene interactions available in BioGrid 3.5.170 [52] compiled on February 25, 2019 as known structural constraints and estimated the weights from data. In the metabolomic data example, we took the metabolic network from KEGG metabolic reactions using the `KEGGgraph` R package (version 1.38.0).

Finally, although NetGSA allows condition-specific networks, we implemented NetGSA assuming equal networks to ensure fair comparisons with topologyGSA and DEGraph.

### Implementation and availability

All methods tested have well-maintained R packages available on CRAN or Bioconductor. Input genes are named by Entrez IDs in all methods with the exception of topologyGSA and CePa, which use instead gene symbols. Pathway topology information was obtained from the KEGG database [37], extracted using the R package graphite on November 28, 2018 for cancer genomic studies, and from KEGG metabolic interactions using the KEGGgraph R package in the metabolomic study.

## Abbreviations

CAMERA: Correlation Adjusted MEan RAnk gene set test
DE: Differentially expressed
DC: Detection call
FCS: Functional class scoring
FDR: False discovery rate
KEGG: Kyoto Encyclopedia of Genes and Genomes
NetGSA: Network-based gene set analysis
ORA: Over-representation analysis
PE: Pathway Express
PRS: Pathway Regulation Score
SPIA: Signaling pathway impact analysis
TCGA: The Cancer Genome Atlas.

## Ethics approval and consent to participate

Not applicable.

## Consent for publication

Not applicable.

## Availability of data and materials

Detailed results for all parameter settings in Excel spreadsheets, together with datasets and R code for reproducing the results of this study can be found in [32].

## Author’s contributions

JM carried out the simulation studies, performed the statistical analyses and wrote the manuscript. AS and GM participated in the design of the simulation studies. AS and GM revised the manuscript and supervised the project. All authors read and approved the final manuscript.

## Acknowledgements

The authors would like to thank three anonymous referees for their constructive comments.

## Figures

### Additional Files

Additional file 1 — Supplementary Materials

Supplementary Materials include details on test design, additional simulation results on the two gene expression data and analysis of a synthetic data set as well as information on data and code availability. (pdf)

