## Supplementary materials for "A Comparative Study of Topology-based Pathway Enrichment Analysis Methods"

*2019-05-05*

**5 Analysis of Metabolomics Data** **12**

**6 Analysis of Synthetic Data** **14**

**References** **16**

### 1 Introduction

In this supplement, we provide details on experimental design and additional numerical results on type I error and power comparisons. We also provide a synthetic data set to illustrate the need for methods that can jointly test for differences in mean expression and network structures.

R code for reproducing numerical results in the paper are available at https://github.com/drjingma/NetGSAreview.

### 2 Test Design

As described in the Experimental Design section, three pathway dysregulation mechanisms were used in the two gene expression data examples, whose characteristics are summarized in Table 1. Specifically, for all topology designs, we set the detection call (DC) to be 10%, meaning that 10% of the genes inside the pathway are differentially expressed with a mean difference varying from 0.1 to 0.5. Note that the magnitude of the mean signal is expressed relative to the unit variance of each gene. Further, at each mean level, we examine the results of enrichment analysis methods with and without permuting the sample labels. When the sample labels are permuted, data from the two experimental conditions share a similar correlation structure. As a result, the significance of each pathway is primarily driven by the differential expression of its members. On the other hand, when the sample labels are fixed to be the same as the ones in the study, it is expected that the two experimental conditions may have different correlation structures. Therefore, enrichment of each pathway could be due to differential expression and/or differences induced due to changes in their interactions.

Table 1: Design of simulation experiments at 10% DC. For each topology design, we consider the original samples and permuted sample labels, and vary the mean signal from 0.1 to 0.5.

| **Topology.design** | Type | Permutation | Mean |
|---|---|---|---|
| **Community** | 1 | no | 0.1 |
| | | | 0.2 |
| | | | 0.3 |
| | | | 0.4 |
| | | | 0.5 |
| | 2 | yes | 0.1 |
| | | | 0.2 |
| | | | 0.3 |
| | | | 0.4 |
| | | | 0.5 |
| **Neighborhood** | 3 | no | 0.1 |
| | | | 0.2 |
| | | | 0.3 |
| | | | 0.4 |
| | | | 0.5 |
| | 4 | yes | 0.1 |
| | | | 0.2 |
| | | | 0.3 |
| | | | 0.4 |
| | | | 0.5 |
| **Betweenness** | 5 | no | 0.1 |
| | | | 0.2 |
| | | | 0.3 |
| | | | 0.4 |
| | | | 0.5 |
| | 6 | yes | 0.1 |
| | | | 0.2 |
| | | | 0.3 |
| | | | 0.4 |
| | | | 0.5 |

### 3 Analysis of Breast Cancer Data

#### 3.1 Type I error

We have already presented the type I errors of 11 KEGG pathways (primarily signaling pathways) in the main article. These pathways are

```
##  [1] "Central carbon metabolism in cancer"
##  [2] "Choline metabolism in cancer"
##  [3] "Sphingolipid signaling pathway"
##  [4] "mTOR signaling pathway"
##  [5] "Adrenergic signaling in cardiomyocytes"
##  [6] "VEGF signaling pathway"
##  [7] "Apelin signaling pathway"
##  [8] "TNF signaling pathway"
##  [9] "Retrograde endocannabinoid signaling"
## [10] "GnRH signaling pathway"
```

```
## [11] "Oxytocin signaling pathway"
```

Additionally, we also look at the type I errors for the following 38 KEGG metabolic pathways:

```
##  [1] "Fructose and mannose metabolism"
##  [2] "Purine metabolism"
##  [3] "Pyrimidine metabolism"
##  [4] "Alanine, aspartate and glutamate metabolism"
##  [5] "Cysteine and methionine metabolism"
##  [6] "Phenylalanine metabolism"
##  [7] "Tryptophan metabolism"
##  [8] "beta-Alanine metabolism"
##  [9] "Selenocompound metabolism"
## [10] "Glutathione metabolism"
## [11] "Starch and sucrose metabolism"
## [12] "Amino sugar and nucleotide sugar metabolism"
## [13] "Inositol phosphate metabolism"
## [14] "Linoleic acid metabolism"
## [15] "alpha-Linolenic acid metabolism"
## [16] "Pyruvate metabolism"
## [17] "Glyoxylate and dicarboxylate metabolism"
## [18] "Propanoate metabolism"
## [19] "Butanoate metabolism"
## [20] "Thiamine metabolism"
## [21] "Riboflavin metabolism"
## [22] "Vitamin B6 metabolism"
## [23] "Nicotinate and nicotinamide metabolism"
## [24] "Porphyrin and chlorophyll metabolism"
## [25] "Sulfur metabolism"
## [26] "Drug metabolism - cytochrome P450"
## [27] "Drug metabolism - other enzymes"
## [28] "Fatty acid biosynthesis"
## [29] "Primary bile acid biosynthesis"
## [30] "Ubiquinone and other terpenoid-quinone biosynthesis"
## [31] "Arginine biosynthesis"
## [32] "Phenylalanine, tyrosine and tryptophan biosynthesis"
## [33] "N-Glycan biosynthesis"
## [34] "Glycosaminoglycan biosynthesis - chondroitin sulfate / dermatan sulfate"
## [35] "Glycosphingolipid biosynthesis - lacto and neolacto series"
## [36] "Pantothenate and CoA biosynthesis"
## [37] "Folate biosynthesis"
## [38] "Aminoacyl-tRNA biosynthesis"
```

These pathways are mostly of small size, and do not satisfy input requirements of topologyGSA, SPIA and Pathway-Express. Therefore, we only compare CAMERA, CePa, DEGraph, NetGSA, PathNet and PRS on the metabolic pathways. Note, however, PRS requires the presence of DE genes, and is not applicable for the self-contained null as there are no DE genes.

Figure 1 shows the type I errors for KEGG metabolic pathways. Overall, we observe that all methods except PathNet control type I errors. Specifically, the type I errors from PathNet for the following several pathways are highly inflated:

```
##                                   name type.I.error
## 460                  Purine metabolism        0.100
## 462              Pyrimidine metabolism        0.085
## 482           Linoleic acid metabolism        0.090
```

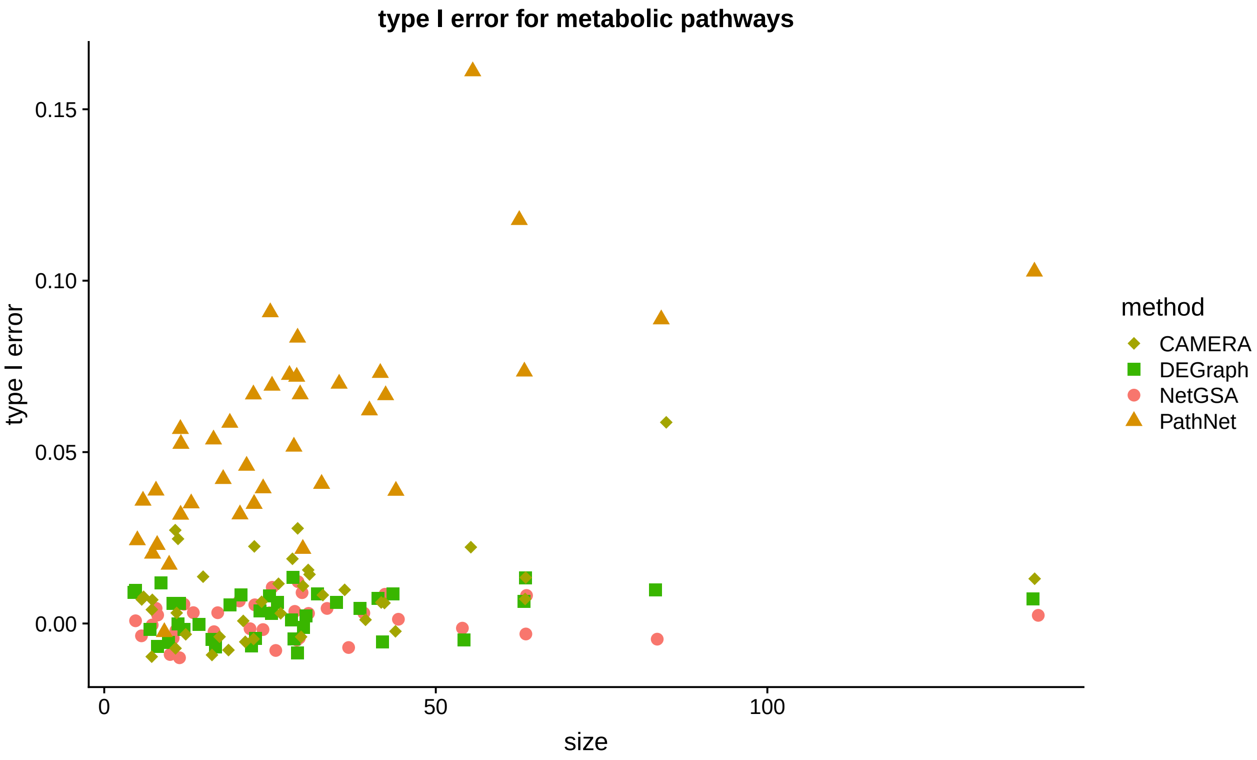

Figure 1: Type I errors for KEGG metabolic pathways in the TCGA breast cancer study (Koboldt et al. 2012). The x-axis shows the pathway size and the y-axis indicates the type I error. Overlapping points are re-positioned by adding ±1 to the x-axis and ±0.01 to the y-axis. Thus the negative values should be understood as being very close to zero. PathNet has inflated type I errors for some pathways, whereas all other methods have slightly conservative type I errors.

```
## 494 Porphyrin and chlorophyll metabolism        0.090
## 496    Drug metabolism - cytochrome P450        0.165
## 497      Drug metabolism - other enzymes        0.110
```

The loss of type I error control with PathNet is likely caused by the inaccurate coverage of gene interactions in these metabolic pathways. For example, the following pathways claim a dense topology (as obtained from the **graphite** package), which may cover many false positive interactions. Because PathNet uses gene-wise fold changes instead of the expression data, these misleading interactions result in inflated type I errors. In contrast, DEGraph and NetGSA combine information from the expression data and pathway topology to achieve control of type I errors.

```
## The graph for Drug metabolism - cytochrome P450 is:
```

```
## A graphNEL graph with directed edges
## Number of Nodes = 55
## Number of Edges = 457
```

```
## The graph for Purine metabolism is:
```

```
## A graphNEL graph with directed edges
## Number of Nodes = 141
## Number of Edges = 6228
```

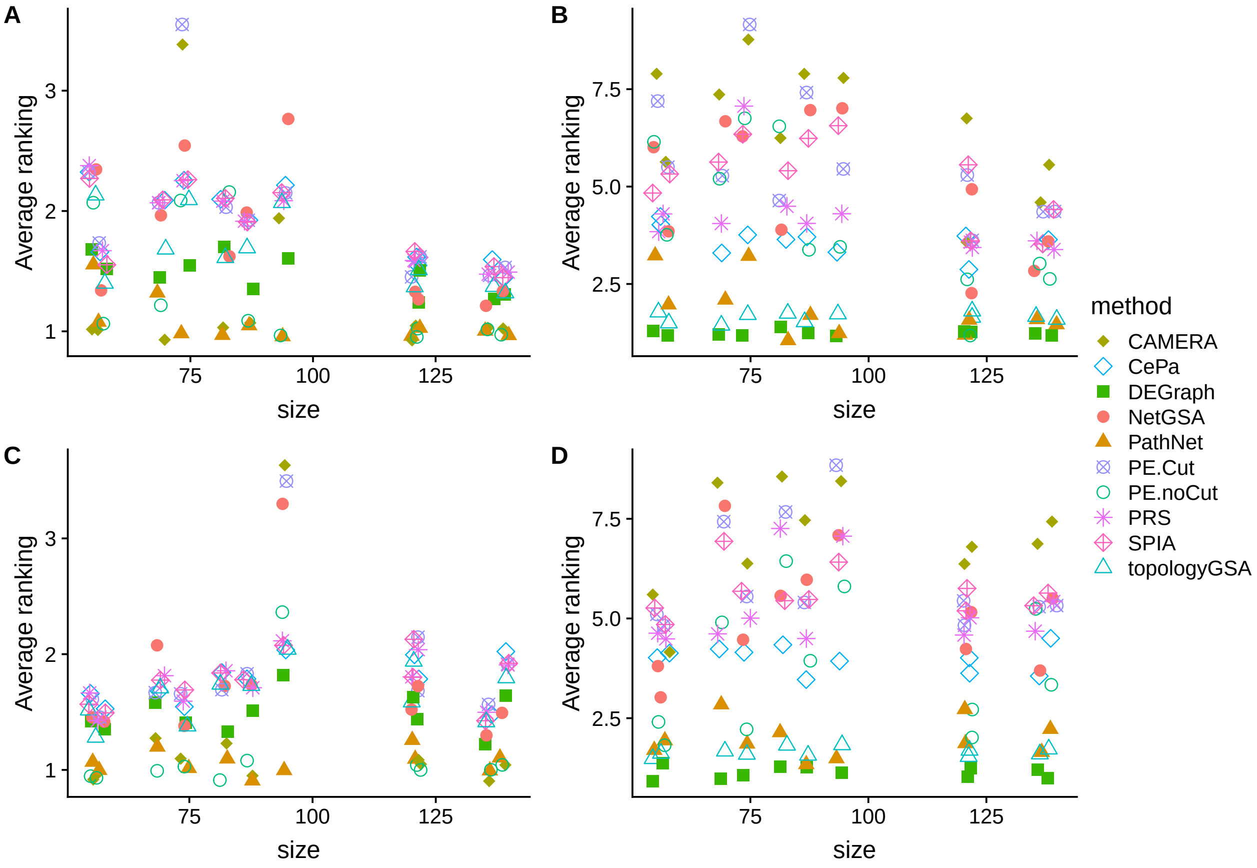

Figure 2: Average ranking of empirical powers on signaling pathways with (A) fixed sample labels and the **community** design; (B) shuffled sample labels and the **community** design; (C) fixed sample labels and the **neighborhood** design; (D) shuffled sample labels and the **neighborhood** design; for the TCGA breast cancer study (Koboldt et al. 2012). The x-axis shows the pathway size, and the y-axis indicates the average ranking of empirical power over different mean changes. Lower ranking indicates better performance. Overlapping points are re-positioned by adding ±1 to the x-axis and ±0.1 to the y-axis.

#### 3.2 Power

As mentioned in the Results section, power comparisons are based on the average ranking of empirical powers across different levels of mean changes. We have already presented the average ranking of the 11 KEGG pathways (primarily signaling pathways) analyzed by all methods under the betweenness dysregulation design in the main article. Below we first show the comparison of average ranking for the 11 pathways under the community and neighborhood dysregulation design, and later show comparisons for KEGG metabolic pathways where only a subset of methods apply.

Figure 2 shows the average ranking of the 11 (primarily signaling) pathways where all methods apply under the **community** and **neighborhood** dysregulation design. The message here is similar to that observed in Figure 3 of the main article. In particular, CAMERA, PathNet and PE.noCut (Pathway-Express without $p$-value cutoff) perform the best when using the original sample labels (Figure 2A, 2C), but DEGraph and topologyGSA outperform the others when using permuted sample labels (Figure 2B, 2D). ORA-type methods rank high (CePa, PE.Cut, PRS and SPIA) because they only work when there are DE genes, i.e. when the mean signal difference is large.

Next we compare the relative power ranking of different methods on metabolic pathways that have at least one DE gene. The set of pathways considered under each dysregulation design thus varies. Specifically, under the **betweenness** design, the following 12 pathways have at least one DE gene:

```
##  [1] "Purine metabolism"
##  [2] "Pyrimidine metabolism"
##  [3] "Alanine, aspartate and glutamate metabolism"
##  [4] "beta-Alanine metabolism"
##  [5] "Starch and sucrose metabolism"
##  [6] "Amino sugar and nucleotide sugar metabolism"
##  [7] "Inositol phosphate metabolism"
##  [8] "Riboflavin metabolism"
##  [9] "Nicotinate and nicotinamide metabolism"
## [10] "Drug metabolism - other enzymes"
## [11] "Arginine biosynthesis"
## [12] "Pantothenate and CoA biosynthesis"
```

Under the **community** design, the following 19 pathways have at least one DE gene:

```
##  [1] "Fructose and mannose metabolism"
##  [2] "Purine metabolism"
##  [3] "Pyrimidine metabolism"
##  [4] "Alanine, aspartate and glutamate metabolism"
##  [5] "Cysteine and methionine metabolism"
##  [6] "Phenylalanine metabolism"
##  [7] "beta-Alanine metabolism"
##  [8] "Starch and sucrose metabolism"
##  [9] "Amino sugar and nucleotide sugar metabolism"
## [10] "Inositol phosphate metabolism"
## [11] "Linoleic acid metabolism"
## [12] "alpha-Linolenic acid metabolism"
## [13] "Pyruvate metabolism"
## [14] "Riboflavin metabolism"
## [15] "Nicotinate and nicotinamide metabolism"
## [16] "Drug metabolism - other enzymes"
## [17] "Arginine biosynthesis"
## [18] "Phenylalanine, tyrosine and tryptophan biosynthesis"
## [19] "Pantothenate and CoA biosynthesis"
```

Under the **neighborhood** design, the following 15 pathways have at least one DE gene:

```
##  [1] "Fructose and mannose metabolism"
##  [2] "Purine metabolism"
##  [3] "Pyrimidine metabolism"
##  [4] "Alanine, aspartate and glutamate metabolism"
##  [5] "Starch and sucrose metabolism"
##  [6] "Amino sugar and nucleotide sugar metabolism"
##  [7] "Inositol phosphate metabolism"
##  [8] "Linoleic acid metabolism"
##  [9] "alpha-Linolenic acid metabolism"
## [10] "Pyruvate metabolism"
## [11] "Nicotinate and nicotinamide metabolism"
## [12] "Drug metabolism - other enzymes"
## [13] "Arginine biosynthesis"
## [14] "Pantothenate and CoA biosynthesis"
## [15] "Folate biosynthesis"
```

Figure 3 shows the average ranking of all six methods under different dysregulation designs. When the sample labels are fixed, no method uniformly dominates the others (Figure 3A, 3C, 3E). CAMERA seems to perform well in several pathways under the **betweenness** design, but perform poorly under the other designs. DEGraph has reliable performances, especially under **community** and **neighborhood** designs (Figure 3D, 3F). On the other hand, with permuted sample labels, DEGraph performs the best (Figure 3B, 3D, 3F). CePa and NetGSA have comparable performances under **community** and **neighborhood** designs (Figure 3D, 3F). The low performance of PRS is partly because it only works when mean signal difference is large.

### 4 Analysis of Prostate Cancer Data

As mentioned in the main article, this data set contains expression levels of 2952 genes across 264 cases and 160 control subjects. We analyzed 112 KEGG signaling and metabolic pathways.

Before power comparison, we first look at type I errors of different methods.

#### 4.1 Type I error

To assess the type I errors, again we look at two sets of pathways. The first subset involves primarily KEGG signaling pathways where all methods apply:

```
##  [1] "Central carbon metabolism in cancer"
##  [2] "Choline metabolism in cancer"
##  [3] "Sphingolipid signaling pathway"
##  [4] "mTOR signaling pathway"
##  [5] "Adrenergic signaling in cardiomyocytes"
##  [6] "VEGF signaling pathway"
##  [7] "Apelin signaling pathway"
##  [8] "TNF signaling pathway"
##  [9] "Retrograde endocannabinoid signaling"
## [10] "GnRH signaling pathway"
## [11] "Oxytocin signaling pathway"
```

and the other set consists of metabolic pathways where only selected methods apply:

```
##  [1] "Fructose and mannose metabolism"
##  [2] "Purine metabolism"
##  [3] "Pyrimidine metabolism"
##  [4] "Alanine, aspartate and glutamate metabolism"
##  [5] "Cysteine and methionine metabolism"
##  [6] "Phenylalanine metabolism"
##  [7] "Tryptophan metabolism"
##  [8] "beta-Alanine metabolism"
##  [9] "Selenocompound metabolism"
## [10] "Glutathione metabolism"
## [11] "Starch and sucrose metabolism"
## [12] "Amino sugar and nucleotide sugar metabolism"
## [13] "Inositol phosphate metabolism"
## [14] "Linoleic acid metabolism"
## [15] "alpha-Linolenic acid metabolism"
## [16] "Pyruvate metabolism"
## [17] "Glyoxylate and dicarboxylate metabolism"
## [18] "Propanoate metabolism"
## [19] "Butanoate metabolism"
## [20] "Thiamine metabolism"
```

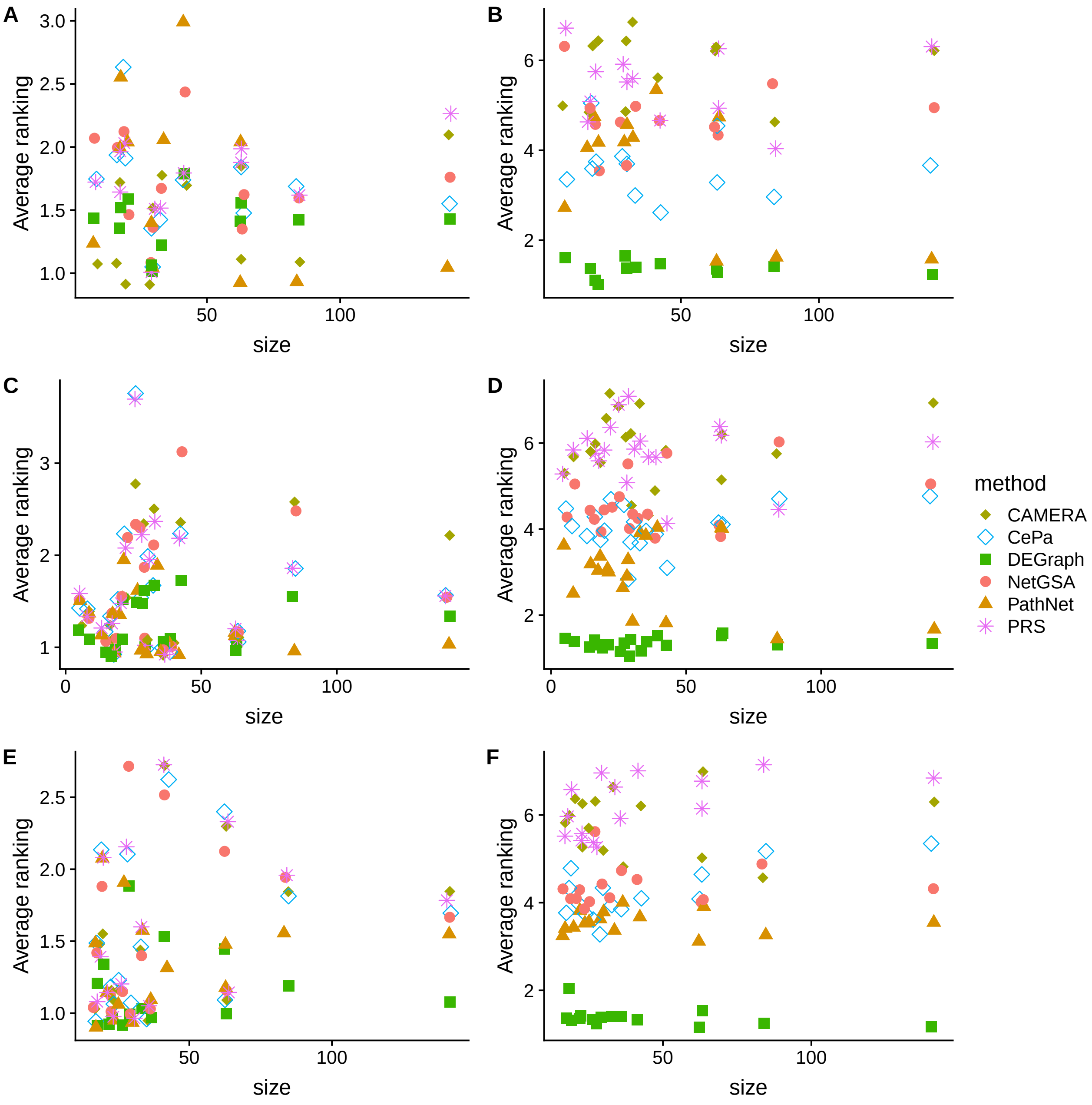

Figure 3: Average ranking of empirical powers on metabolic pathways with (A) fixed sample labels and the **betweenness** design; (B) shuffled sample labels and the **betweenness** design; (C) fixed sample labels and the **community** design; (D) shuffled sample labels and the **community** design; (E) fixed sample labels and the **neighborhood** design; (F) shuffled sample labels and the **neighborhood** design, for the TCGA breast cancer study (Koboldt et al. 2012). The x-axis shows the pathway size, and the y-axis indicates the average ranking of empirical power over different mean changes. Lower ranking indicates better performance. Overlapping points are re-positioned by adding ±1 to the x-axis and ±0.1 to the y-axis. No method uniformly dominates when using the original sample labels (A,C,E), but DEGraph yields the best performance with permuted sample labels (B,D,F).

```
## [21] "Riboflavin metabolism"
## [22] "Nicotinate and nicotinamide metabolism"
## [23] "Porphyrin and chlorophyll metabolism"
## [24] "Sulfur metabolism"
## [25] "Drug metabolism - cytochrome P450"
## [26] "Drug metabolism - other enzymes"
## [27] "Fatty acid biosynthesis"
## [28] "Primary bile acid biosynthesis"
## [29] "Ubiquinone and other terpenoid-quinone biosynthesis"
## [30] "Arginine biosynthesis"
## [31] "N-Glycan biosynthesis"
## [32] "Glycosaminoglycan biosynthesis - chondroitin sulfate / dermatan sulfate"
## [33] "Glycosphingolipid biosynthesis - lacto and neolacto series"
## [34] "Pantothenate and CoA biosynthesis"
## [35] "Folate biosynthesis"
## [36] "Aminoacyl-tRNA biosynthesis"
```

Figure 4 shows the type I errors for the two types of pathways. The message here is similar to that seen in the breast cancer data example, except that PE.noCut exhibits inflated type I errors for several signaling pathways (Figure 4A), which are *mTOR signaling pathway* (type I error 0.15), *GnRH signaling pathway* (type I error 0.125), and *TNF signaling pathway* (type I error 0.09). Type I errors of PathNet on metabolic pathways again are slightly inflated in several instances (Figure 4B), such as *Pyrimidine metabolism* (type I error 0.13) and *Purine metabolism* (type I error 0.16). Again, this could be because both *Pyrimidine metabolism* and *Purine metabolism* retrieved from the **graphite** package consist of many false positives edges:

```
## The graph for Pyrimidine metabolism is:
```

```
## A graphNEL graph with directed edges
## Number of Nodes = 88
## Number of Edges = 1896
```

```
## The graph for Purine metabolism is:
```

```
## A graphNEL graph with directed edges
## Number of Nodes = 153
## Number of Edges = 6923
```

#### 4.2 Power

As discussed in the previous section, power comparisons are based on the average ranking of empirical powers across different levels of mean changes. Below we first show the comparison of average ranking for signaling pathways where all methods apply, and later show comparisons for metabolic pathways where only a subset of methods apply.

Figure 5 shows the relative performance of different methods on the 11 (primarily signaling) pathways where all methods apply under different dysregulation designs. When sample labels are fixed, CAMERA, PathNet and PE.noCut rank the best (Figure 5A, 5C), though CAMERA falls short under the neighborhood design (Figure 5E). With shuffled sample labels, DEGraph, PathNet and topologyGSA perform the best (Figure 5B,5D,5F).

Next we compare the relative power ranking of different methods on metabolic pathways that have at least one DE gene. The set of pathways considered under each dysregulation design thus varies. Specifically, under the **betweenness** design, the following 18 pathways have at least one DE gene:

```
##  [1] "Purine metabolism"
##  [2] "Pyrimidine metabolism"
##  [3] "Alanine, aspartate and glutamate metabolism"
```

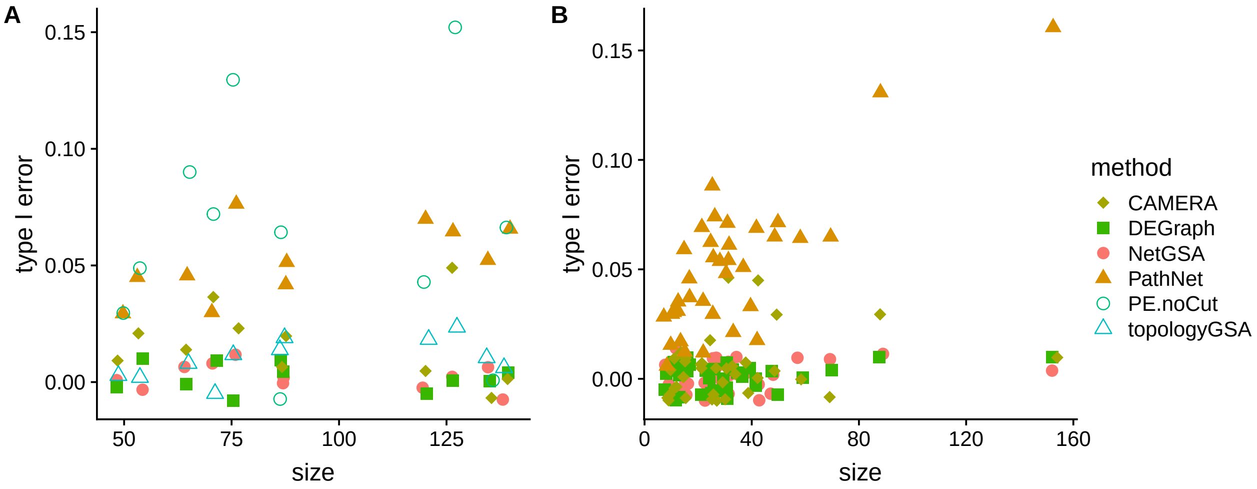

Figure 4: Type I errors for KEGG signaling (A) and metabolic (B) pathways in the TCGA prostate cancer study (Abeshouse et al. 2015). The x-axis shows the pathway size and the y-axis indicates the type I error. Overlapping points are re-positioned by adding ±1 to the x-axis and ±0.01 to the y-axis. Thus the negative values should be understood as being very close to zero.

```
##  [4] "Phenylalanine metabolism"
##  [5] "Tryptophan metabolism"
##  [6] "Starch and sucrose metabolism"
##  [7] "Amino sugar and nucleotide sugar metabolism"
##  [8] "Inositol phosphate metabolism"
##  [9] "Pyruvate metabolism"
## [10] "Glyoxylate and dicarboxylate metabolism"
## [11] "Riboflavin metabolism"
## [12] "Nicotinate and nicotinamide metabolism"
## [13] "Drug metabolism - cytochrome P450"
## [14] "Drug metabolism - other enzymes"
## [15] "Ubiquinone and other terpenoid-quinone biosynthesis"
## [16] "Arginine biosynthesis"
## [17] "Pantothenate and CoA biosynthesis"
## [18] "Aminoacyl-tRNA biosynthesis"
```

Under the **community** design, the following 19 pathways have at least one DE gene:

```
##  [1] "Fructose and mannose metabolism"
##  [2] "Purine metabolism"
##  [3] "Pyrimidine metabolism"
##  [4] "Alanine, aspartate and glutamate metabolism"
##  [5] "Cysteine and methionine metabolism"
##  [6] "Phenylalanine metabolism"
##  [7] "Tryptophan metabolism"
##  [8] "beta-Alanine metabolism"
##  [9] "Starch and sucrose metabolism"
## [10] "Amino sugar and nucleotide sugar metabolism"
## [11] "Inositol phosphate metabolism"
## [12] "Linoleic acid metabolism"
## [13] "alpha-Linolenic acid metabolism"
## [14] "Glyoxylate and dicarboxylate metabolism"
```

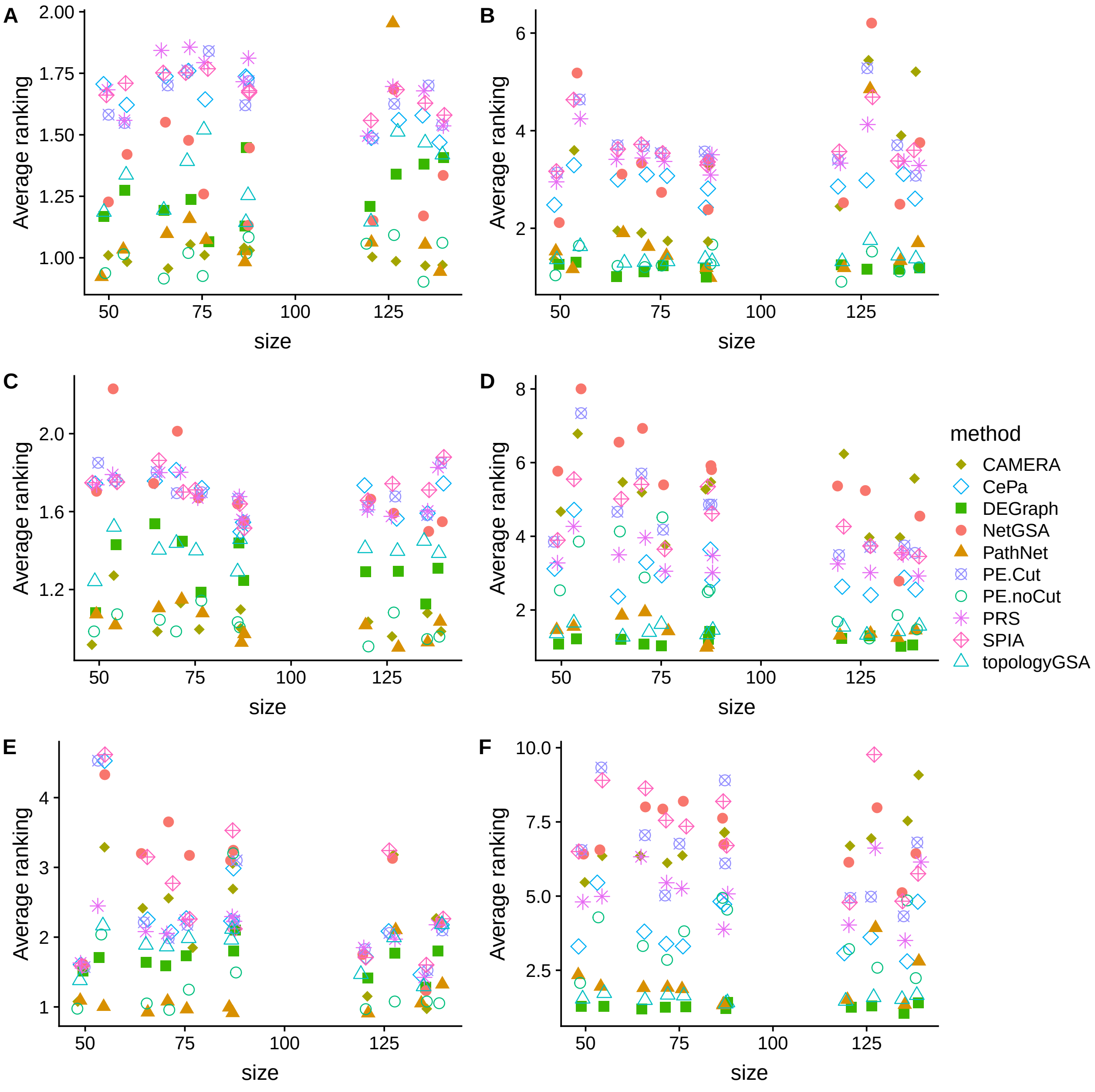

Figure 5: Average ranking of empirical powers on signaling pathways with (A) fixed sample labels and the **betweenness** design; (B) shuffled sample labels and the **betweenness** design; (C) fixed sample labels and the **community** design; (D) shuffled sample labels and the **community** design; (E) fixed sample labels and the **neighborhood** design; (F) shuffled sample labels and the **neighborhood** design; for the TCGA prostate cancer study (Abeshouse et al. 2015). The x-axis shows the pathway size, and the y-axis indicates the average ranking of empirical power over different mean changes. Lower ranking indicates better performance. Overlapping points are re-positioned by adding ±1 to the x-axis and ±0.1 to the y-axis.

```
## [15] "Nicotinate and nicotinamide metabolism"
## [16] "Drug metabolism - cytochrome P450"
## [17] "Drug metabolism - other enzymes"
## [18] "Arginine biosynthesis"
## [19] "Aminoacyl-tRNA biosynthesis"
```

Under the **neighborhood** design, the following 16 pathways have at least one DE gene:

```
##  [1] "Fructose and mannose metabolism"
##  [2] "Purine metabolism"
##  [3] "Pyrimidine metabolism"
##  [4] "Alanine, aspartate and glutamate metabolism"
##  [5] "Phenylalanine metabolism"
##  [6] "beta-Alanine metabolism"
##  [7] "Starch and sucrose metabolism"
##  [8] "Amino sugar and nucleotide sugar metabolism"
##  [9] "Inositol phosphate metabolism"
## [10] "Glyoxylate and dicarboxylate metabolism"
## [11] "Nicotinate and nicotinamide metabolism"
## [12] "Porphyrin and chlorophyll metabolism"
## [13] "Drug metabolism - cytochrome P450"
## [14] "Drug metabolism - other enzymes"
## [15] "Arginine biosynthesis"
## [16] "Aminoacyl-tRNA biosynthesis"
```

Figure 6 shows the relative performance of different methods when analyzing metabolic pathways. When the sample labels are fixed, no method uniformly dominates the others (Figure 6A, 6C, 6E). DEGraph seems to yield the most reliable performance throughout. On the other hand, with permuted sample labels, DEGraph performs the best (Figure 6B, 6D, 6F). CePa and NetGSA have comparable performances under **neighborhood** design (Figure 6F). Methods that test competitive null such as CAMERA and PRS perform poorly when sample labels are shuffled.

### 5 Analysis of Metabolomics Data

The metabolomics data example considers the following metabolic pathways:

```
##  [1] "2-Oxocarboxylic acid metabolism "
##  [2] "Alanine, aspartate and glutamate metabolism "
##  [3] "Aminoacyl-tRNA biosynthesis "
##  [4] "Arginine and proline metabolism "
##  [5] "Carbohydrate digestion and absorption "
##  [6] "Citrate cycle (TCA cycle) "
##  [7] "Cysteine and methionine metabolism "
##  [8] "Drug metabolism - other enzymes "
##  [9] "Fructose and mannose metabolism "
## [10] "GABAergic synapse "
## [11] "Galactose metabolism "
## [12] "Glutathione metabolism "
## [13] "Glycerolipid metabolism "
## [14] "Glycine, serine and threonine metabolism "
## [15] "Glycosphingolipid biosynthesis - globo series "
## [16] "Glyoxylate and dicarboxylate metabolism "
## [17] "Histidine metabolism "
## [18] "Lysine biosynthesis "
```

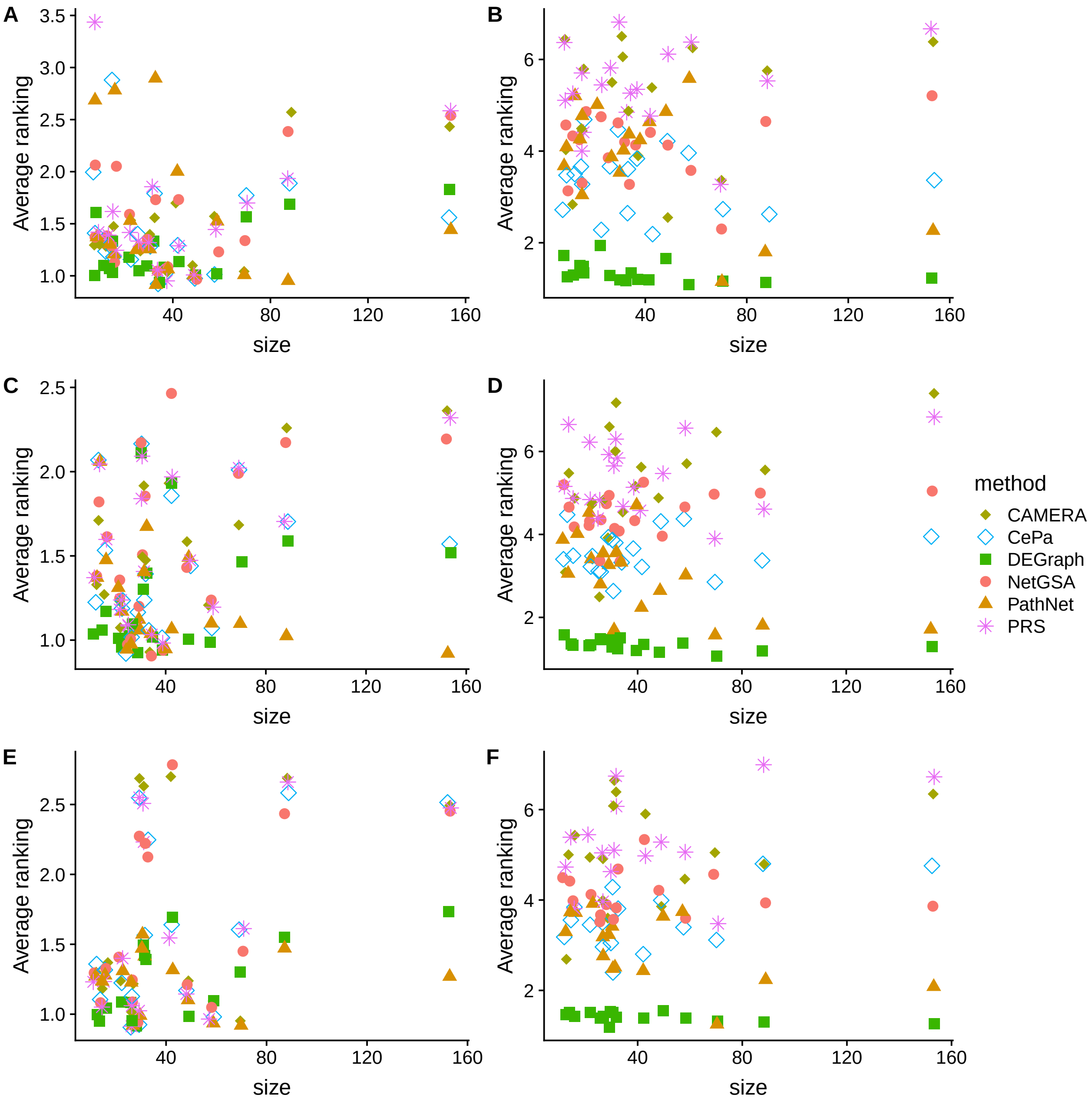

Figure 6: Average ranking of empirical powers on metabolic pathways with (A) fixed sample labels and the **betweenness** design; (B) shuffled sample labels and the **betweenness** design; (C) fixed sample labels and the **community** design; (D) shuffled sample labels and the **community** design; (E) fixed sample labels and the **neighborhood** design; (F) shuffled sample labels and the **neighborhood** design, for the TCGA prostate cancer study (Abeshouse et al. 2015). The x-axis shows the pathway size, and the y-axis indicates the average ranking of empirical power over different mean changes. Lower ranking indicates better performance. Overlapping points are re-positioned by adding ±1 to the x-axis and ±0.1 to the y-axis. No method uniformly dominates when using the original sample labels (A,C,E), but DEGraph yields the best performance with permuted sample labels (B,D,F).

```
## [19] "Lysosome "
## [20] "Nicotinate and nicotinamide metabolism "
## [21] "Pantothenate and CoA biosynthesis "
## [22] "Peroxisome "
## [23] "Phenylalanine metabolism "
## [24] "Purine metabolism "
## [25] "Pyrimidine metabolism "
## [26] "Pyruvate metabolism "
## [27] "Selenocompound metabolism "
## [28] "Sphingolipid metabolism "
## [29] "Starch and sucrose metabolism "
## [30] "Tryptophan metabolism "
## [31] "Tyrosine metabolism "
## [32] "Valine, leucine and isoleucine degradation "
```

These KEGG metabolic pathways are mostly of small size, and tend to overlap. Figure 7 illustrates how many members are shared among pathways. Black cell indicates presence of a metabolite in the corresponding pathway and white indicates absence. Most noticebly, *Alanine, aspartate and glutamate metabolism* (size 23) and *Aminoacyl-tRNA biosynthesis* (size 20) have 15 metabolites in common, which constitute a majority of their members.

### 6 Analysis of Synthetic Data

Lastly, we present a simulation example based on synthetic data to illustrate the advantage of NetGSA in detecting both changes in mean expression levels and/or changes in network structure. Figure 8 shows the network and subnetworks being compared. A striking feature is

As indicated by node shapes, subnetworks 1 and 6 have equal means under the two conditions, whereas subnetworks 3 and 8, 4 and 5, 2 and 7 have, respectively, 20%, 40% and 60% nodes with differential means. Further, some subnetworks also have changes in network topology. In particular, the topologies of subnetworks 4, 6, 7 and 8 remain the same under both conditions, whereas those of subnetworks 1, 2, 3, and 5 are completely rewired. Such dramatic changes in network topology are often expected in practice. In this case, DEGraph is not recommended because it assumes the network structures under the two conditions are the same, when in fact they are not. Whichever network provided for DEGraph will misspecify the graph topology for at least one condition. In contrast, NetGSA can leverage both changes in mean expressions and changes in network structures.

The empirical powers for each subnetwork averaged in 100 replications are shown in Table 2. Here the results for NetGSA were obtained assuming the network under each condition is known *a priori*. The network provided for DEGraph is the network under the null. As expected, NetGSA identifies 5 (0.99) and 3 (0.96) as the most significantly enriched pathways due to both changes in mean and in network topology, followed by pathway 7 (0.94) and 2 (0.89). An interesting observation is that NetGSA assigns slightly higher power for pathway 7 compared to pathway 2. This is because some of the DE nodes in pathway 7 are in *hub* positions. In comparison, DEGraph correctly identified pathway 2, 3, and 5 as the enriched ones, but missed pathway 7.

It is important to note that it is unfair to compare NetGSA and DEGraph using this synthetic data example when the network for DEGraph has to be misspecified. Nonetheless, this example illustrates how NetGSA can be used to simultaneously detect changes in mean and network structures.

Table 2: Empirical powers averaged in 100 replications with 50 observations per condition

| Method | set.1 | set.2 | set.3 | set.4 | set.5 | set.6 | set.7 | set.8 |
|---|---|---|---|---|---|---|---|---|
| NetGSA | 0.08 | 0.89 | 0.96 | 0.14 | 0.99 | 0.02 | 0.94 | 0.03 |
| DEGraph | 0.18 | 1.00 | 1.00 | 0.49 | 1.00 | 0.06 | 0.62 | 0.31 |
| true power | 0.12 | 0.93 | 0.98 | 0.11 | 0.99 | 0.05 | 0.95 | 0.10 |

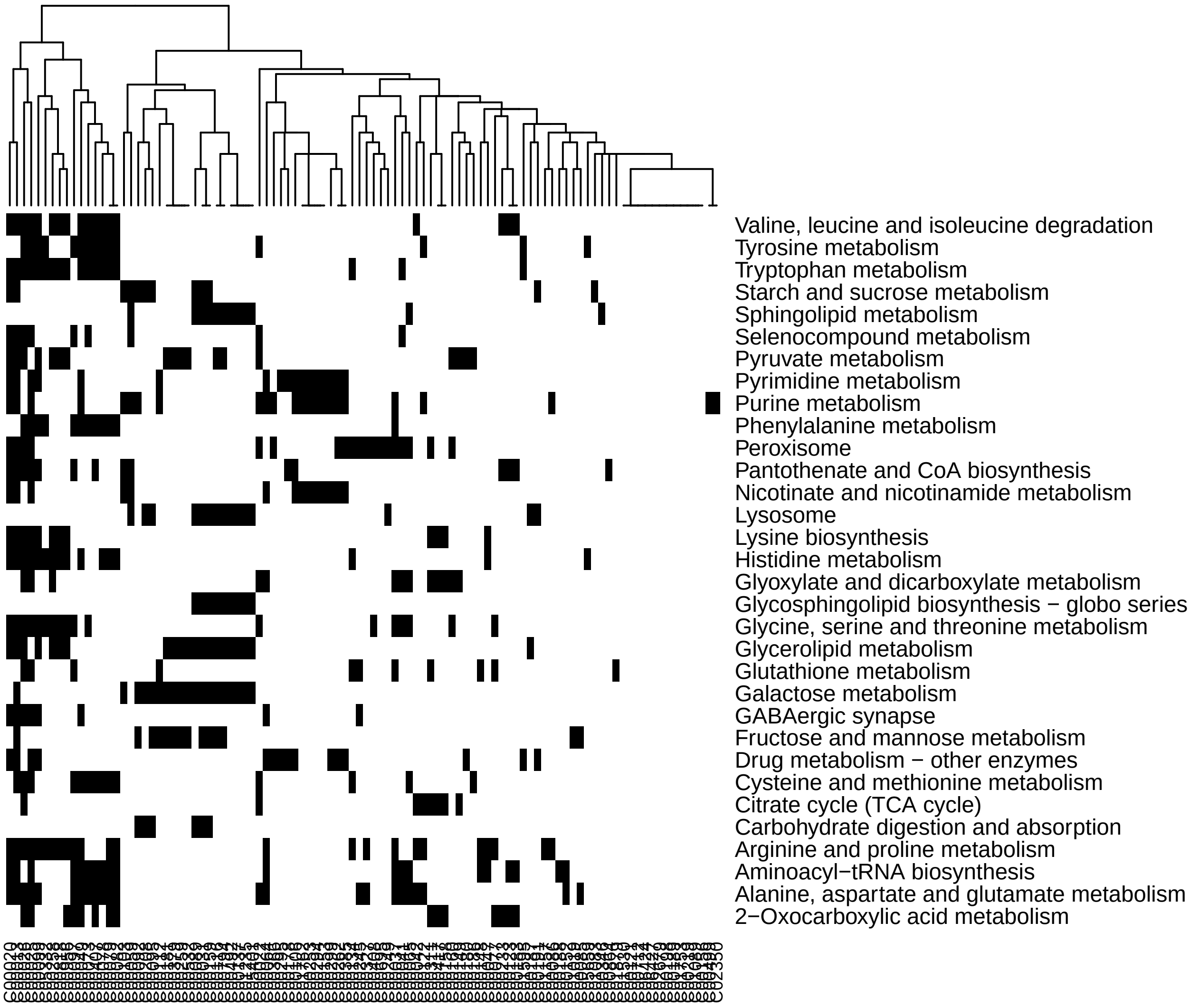

Figure 7: Heatmap of pathway membership in the metabolomics data example

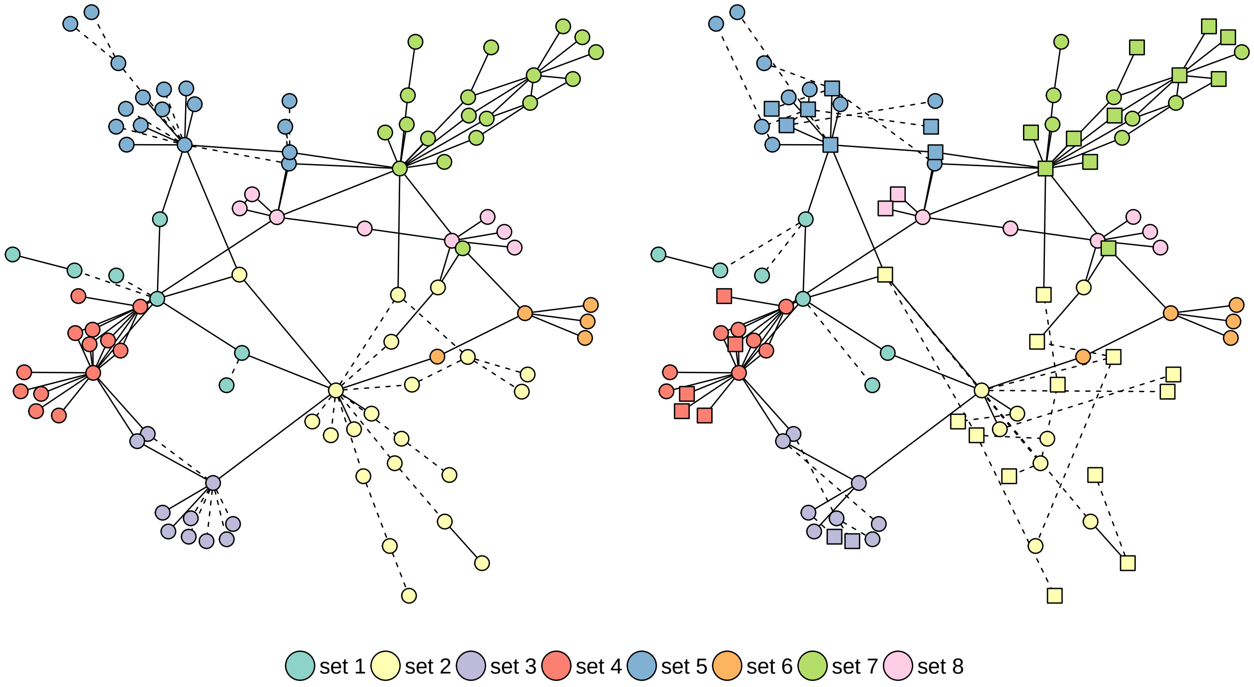

Figure 8: The network and subnetwork topology under the null (left) and alternative (right). Dashed lines represent edges that are present in only one condition. Nodes in square are associated with mean changes.
